# KAT6 and KAT7 actively maintain mitotic chromatin composition and acetylation to preserve developmental competence

**DOI:** 10.64898/2026.11.07.765049

**Authors:** Alejandra Laguillo-Diego, UkJin Lee, Xiao Liu, Abigail Lidoski, Kritika Kasliwal, Urko del Castillo, Jin Joo Kang, Stephanie Stransky, Spencer Kaufman, Bryce J. Katch, Paraskevi Giannakakou, Simone Sidoli, Haiyuan Yu, Matthias Stadtfeld, Effie Apostolou

**Author notes:** Corresponding author: Effie Apostolou.

## Abstract

Mitotic bookmarking enables transmission of regulatory information across cell division, yet how active chromatin states are molecularly preserved during mitosis and how they contribute to postmitotic cell fate decisions remain unresolved. Here, we demonstrate that the acetyltransferases KAT6 and KAT7 remain chromatin-bound and catalytically active throughout mitosis in mouse and human pluripotent stem cells. Acute KAT6/7 depletion disrupts mitotic histone acetylation and the retention of multiple chromatin regulators with minimal effects on chromatin accessibility. Mechanistically, KAT6/7 bookmark both self-renewal and developmental gene loci, where they are required for post-mitotic transcriptional reactivation and priming of lineage-specific programs. Consistently, genes impaired during postmitotic reactivation also fail to activate during neuronal differentiation upon continuous KAT6/7 perturbation, compromising the establishment of the neuronal program and identity. Altogether, our findings identify KAT6 and KAT7 as a cooperative mitotic chromatin maintenance system that couples active histone acetylation to preservation of chromatin composition, transcriptional memory, and developmental competence across cell division, extending the bookmarking function beyond transcriptional memory. These findings also provide a new framework for understanding KAT6/7 function in neurodevelopmental disorders and cancer.

## Introduction

Pluripotent stem cells are endowed with the unique ability to self-renew indefinitely while retaining the capacity to generate all embryonic lineages^1,2^. The decision to either maintain or change their identity is closely linked to the transition from mitosis to G1 phase^3–5^, when cells emerge from a period of global transcriptional silencing, chromatin compaction, and widespread displacement of chromatin-associated factors^6–9^. As transcription resumes, stem cells must rapidly re-establish self-renewal programs while remaining competent to respond to developmental cues. Accordingly, several lineage regulators are transiently activated during early G1, suggesting that post-mitotic transcriptional resetting contributes to lineage priming^10–12^. However, the molecular mechanisms that coordinate preservation of stem-cell identity with activation of differentiation programs during the mitosis-to-G1 transition remain poorly understood.

One mechanism proposed to mediate the transmission of regulatory information through cell division is mitotic bookmarking, the selective retention of proteins and chromatin features on mitotic chromosomes that facilitate post-mitotic transcriptional reactivation^8,13,14^. Mitotic bookmarking has been reported for several transcription factors, chromatin modifiers remodelers, chromatin modifiers and oncogenic fusion proteins in various contexts^13–15^. Similarly, histone methylation marks associated both with gene silencing (e.g. H3K27me3 and H3K9me3) or activation (e.g, H3K4me3) were shown to be broadly maintained through cell division^8,16–21^. In contrast, the role of histone acetylation in epigenetic memory remains controversial. Histone acetylation is generally associated with active transcription and accessible chromatin and has long been thought to be largely erased during mitosis^8,22–24^. However, several studies reported partial persistence of histone acetylation on mitotic chromatin^17,25–29^, including H3K27ac bookmarking of cell identity genes and enhancers^32,33^, while recent quantitative proteomic analyses documented widespread retention across diverse cell type^30,31^. Importantly, temporal perturbation of H3K27ac during mitosis delayed post-mitotic transcriptional reactivation^12^, arguing for a functional role of mitotic acetylation in the transmission of active chromatin states and transcriptional memory.

The rapid turnover of histone acetylation, relative to histone methylation, has long been used as an argument against its role as a carrier of epigenetic memory. More than two decades ago, it was proposed that the persistence of acetylation marks during mitosis will likely require continued histone acetyltransferase (HAT) activity^34^, although direct evidence for this model was lacking. While several studies have documented the presence and activity of histone deacetylases (HDACs) during mitosis^35–37^, the behavior and function of HATs remain far less understood. Conflicting reports have suggested either widespread eviction of HATs from mitotic chromatin^22,38^ or selective retention of individual enzymes on mitotic chromosomes (i.e: EP300, HAT1, KAT7)^36–39^. Therefore, it is yet unclear to what degree HATs remain bound and catalytically active on mitotic chromatin, and -more importantly-whether their activity contributes to chromatin bookmarking, epigenetic memory, or post-mitotic cell fate decisions.

Here, using a combination of unbiased mass spectrometry, live-cell imaging and/or Chromatin Immunoprecipitation, we provide evidence of a widespread presence of HATs and acetylation marks on mitotic chromatin both in mouse and human ESCs. We focus on KAT7 and KAT6A/B, two HATs that operate in multiprotein complexes and share common subunits and target acetylation marks^40^. These HATs have increasingly appreciated roles in cancer and neurodevelopmental disorders^41–50^, but unclear functional synergies and mechanistic roles in mitotic inheritance. Mechanistically, we show that these enzymes bind specific regions during mitosis and actively maintain mitotic histone acetylation in a cooperative fashion, while they also preserve mitotic chromatin composition, including retention of other bookmarks. In mouse ESCs, KAT7 and KAT6B jointly regulate the transcriptional resetting of stem cell-related genes but also the priming of lineage-specific genes, such as those related to neuronal differentiation. Extending these findings to human ESC, we demonstrate that KAT6/7 activity is required for efficient activation of neurodevelopmental gene networks and neuronal differentiation. Together, our findings reveal that mitotic chromatin acetylation and composition are actively maintained, and uncover a dual role for KAT6/7-dependent bookmarking in preserving cellular identity, while transmitting developmental competence across cell division.

## Results

### Histone acetylation marks are widely retained on mitotic chromatin in mESCs

To determine the retention of histone acetylation marks during mitosis in pluripotent cells, we performed unbiased quantitative histone mass spectrometry (histone MS) on both mitotic and asynchronous mouse embryonic stem cells (mESCs). To do so, we obtained highly pure mitotic cells by synchronization with a dual CDK1 inhibitor/nocodazole treatment (Figure 1A), as assayed by FACS analysis for DNA content and by immunofluorescence for the mitotic marker Histone 3 phospho-Serine 10 (H3S10ph) (Suppl. Figure 1A-B). The relative abundance of each histone modification was calculated as previously described^51^ and compared between mitotic and asynchronous samples (Suppl. Table 1). Consistent with a high mitotic index, H3 phosphorylation (H3S10Ph) was significantly enriched in the mitotic samples (Figure 1B). On the other hand, the global H3me and H4me levels, as well as the individual histone methylation marks, were largely retained in mitotic cells, in agreement with previous reports^16,19–22^ (Figure 1B and Suppl. Figure 1C). Similarly, global H3ac levels and individual H3ac modifications remained largely unchanged during mitosis, whereas several H4ac peptides showed a modest, but significant reduction (Figure 1B-C). These trends were independently validated by Western blot (WB) analysis and immunofluorescence imaging, confirming a preferential retention of the H3ac peptides over H4ac (Figure 1D-E and Suppl. Figure 1E). Although H3K27ac and H3K9ac were not detected in our unbiased histone MS due to their low abundance in the genome, WB analysis confirmed their substantial retention on mitotic chromatin, in agreement with previous publications^32^ (Suppl. Figure 1D). Altogether, these orthogonal approaches demonstrate that histone acetylation is broadly preserved during mitosis, in agreement with recent studies in mESCs and other cell types^30–32^, with H3 acetylation showing greater retention than H4 acetylation.

**Figure 1.**
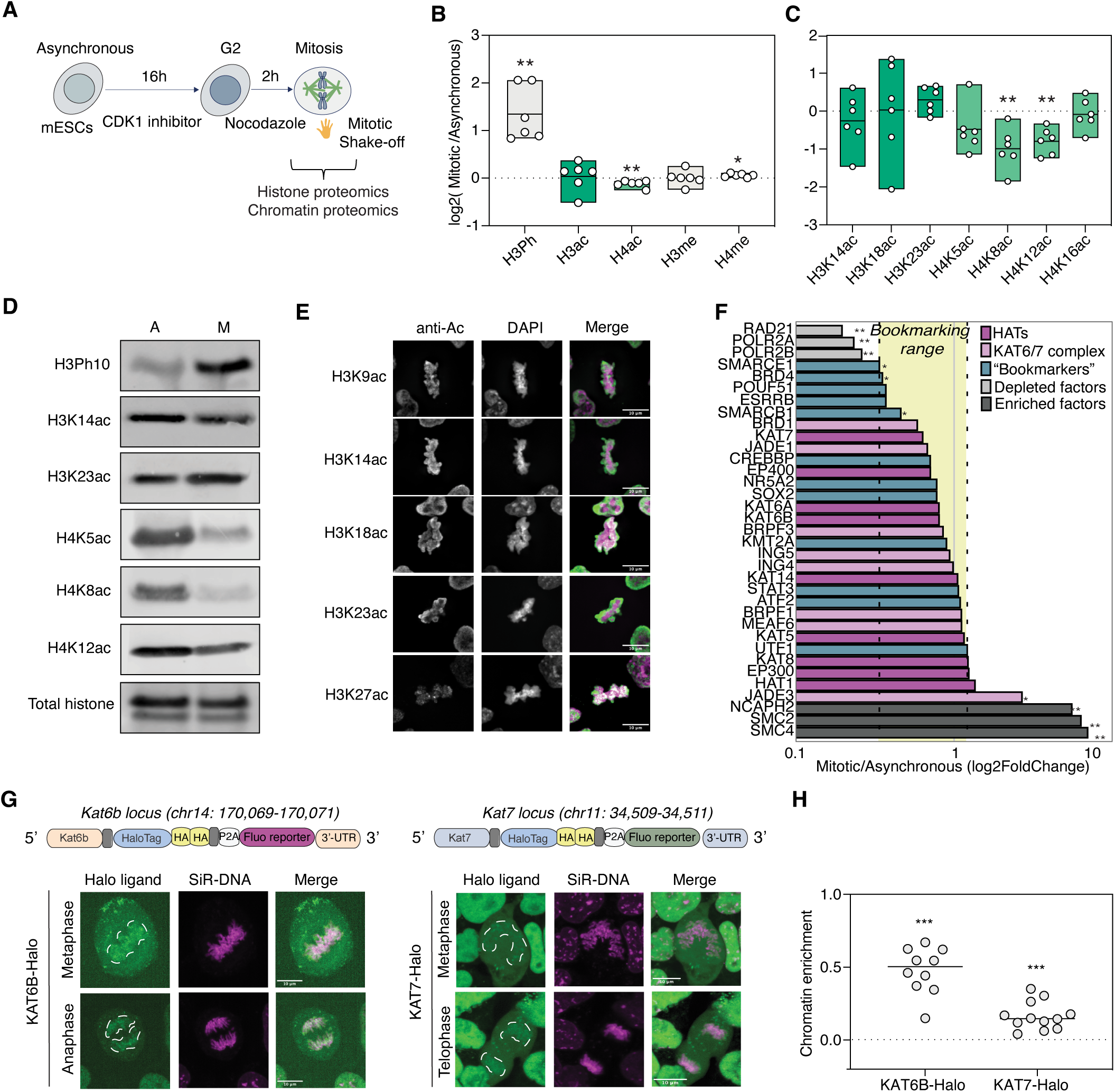
Widespread retention of histone acetylation marks and histone acetyltransferases (HATs) on mitotic chromatin in mESCs. **A.** Experimental strategy for collection and characterization of mitotic and asynchronous mESCs by mass-spectrometry analysis of histone and chromatin-bound extracts. ESCs were synchronized in mitosis by a dual-CDK1 inhibitor/nocodazole treatment, collected by mitotic shake-off and subjected to histone and chromatin proteomics. **B.** Box plot comparing the relative abundance of H3Ph, H3ac, H4ac, H3me and H4me between mitotic and asynchronous extracts as log2 ratio. Data from n = 6 biological replicates are shown. Statistical significance was assessed by one-sample two-tailed t-test against 0. *p < 0.05, **p < 0.01. **C.** Box plot comparing the relative abundance across select H3ac residues (dark green) and H4ac residues (lighter green) between mitotic and asynchronous extracts as log2 ratio. Data from n = 6 biological replicates are shown. Statistical significance was assessed by one-sample two-tailed t-test against 0. **p < 0.01. The full histone proteomics data is provided in Suppl. Table 1. **D.** Western blot analysis showing asynchronous and mitotic levels of select H3ac and H4ac histone marks. Total histone was used as a loading control. Representative of n = 2 biological replicates. **E.** Immunofluorescence analysis for H3ac marks. Representative images of metaphasic chromatin staining for H39ac, H3K14ac, H3K18ac, H3K23ac and H3K27ac are shown. Acetylated histones and DAPI are depicted in gray individually and in green and magenta respectively in the merge. Scale bar = 10 μm. **F.** Bar plot of the mitotic/asynchronous ranking, log2foldchange(mitotic/asynchronous), in logarithmic scale for different categories of proteins. Proteins known to be either mitotically enriched or depleted are shown as controls in grey. Previously described bookmarking factors in mESCs are shown in blue and are used to set the “bookmarking range” (yellow shadow limited by two dotted lines). Histone acetyltransferases (HATs), pink, and KAT6/7 complex subunits, violet, are present within the defined “bookmarking range”. Differential analysis was performed using MAGMa^81^ on quadruplets (2 biological replicates including 2 technical replicates each) statistics are shown as *p < 0.05, **p < 0.01. The full analysis is provided in Suppl. Table 2. **G.** *Top:* schematic of the HaloTag® CRISPR knock-in strategy in mouse ESCs to generate endogenous, fusion KAT6B-Halo and KAT7-Halo proteins for ligand-mediated live imaging. The knock-in cassettes were introduced into the endogenous Kat7 and Kat6b gene *loci* after the last exon. In addition to the HaloTag® tag surrounded by linkers, the cassette contains 2xHA tags and a fluorescent transcriptional reporter preceded by P2A tag to facilitate cell-line generation. *Bottom:* live-cell imaging of mESCs HalotTagged for KAT6B and KAT7. Representative images show the HaloTagged protein with a dotted white line circling the chromatin in different mitotic phases. DNA is shown in magenta. Scale bar = 10 μm. **H.** Dot plot showing the chromatin enrichment (Chr. Enr) for both HaloTagged proteins calculated as log_2_(Chromosome intensity/Cytoplasmic intensity). Chr. Enr > 0.0 indicates mitotic enrichment while Chr. Enr < 0.0 shows relative depletion from mitotic chromosomes. n = 10 cells for mKAT6B, n = 12 cells for mKAT7. Both proteins were significantly enriched on mitotic chromatin as assessed by one-sample two-tailed t-test (***p< 0.001).

### The histone acetyltransferases KAT6B and KAT7 are partially retained on mitotic chromatin in mESCs

As histone acetylation has a short half-life compared to methylation^52^, our observed widespread retention of mitotic histone acetylation implies an active deposition of histone acetylation marks by histone acetyltransferases (HATs) during M-phase. Yet, the association of HATs with mitotic chromatin remains controversial^22,36,38^. To systematically determine the degree of HATs’ retention on mitotic chromatin, we performed subcellular fractionation in asynchronous and mitotic ESCs (Figure 1A) followed by quantitative mass spectrometry of their chromatin-bound fractions. We found 405 proteins to be significantly depleted from mitotic chromatin, while 124 were enriched (Suppl. Figure 1F and Suppl. Table 2). In agreement with the transcriptional silencing during mitosis^53^, RNA polymerase subunits (i.e., POLR2A, POLR2B) and splicing factors (i.e., SF3B1, PRPF8) were strongly depleted from mitotic extracts (Figure 1F). On the other hand, condensin subunits (i.e., SMC4, SMC2, NCAPH, NCAPH2, NCAPD3) were highly enriched in the mitotic chromatin fractions, concordant with their role in mitotic chromosome condensation (Figure 1F). Most proteins were in the ‘grey area’, neither significantly depleted nor enriched, indicating partial conservation of the ‘chromatome’ during mitosis (Suppl. Figure 1F). As expected, many of the previously described mitotic bookmarking factors in mESCs ^14,54^ fell into this grey zone (Figure 1F and Suppl. Figure 1F).

Surprisingly, all major HATs were also found to be preserved on mitotic chromatin to the same degree as previously reported bookmarks (Figure 1F and Suppl. Figure 1F), providing a molecular basis for the observed retention of histone acetylation marks. Among HATs that target the top-retained H3ac residues, H3K23ac and H3K14ac, the paralogous KAT6A/B (MOZ/MORF; H3K23ac and H3K14ac)^40^, and KAT7 (HBO1/MYST; H3K14ac)^40^, were consistently detected as mitotically retained, in agreement with a recent proteomics study on purified metaphase chromosomes in mESCs^55^ (Suppl. Figure 1G). Interestingly, many of the core components of the MOZ/MORF and HBO1 complexes (ie., MEAF6, ING4/5), were also partially preserved during mitosis (Figure 1F), indicating that the presence of catalytically active complex on mitotic chromatin.

To directly visualize candidate HATs’ localization throughout mitosis, we engineered CRISPR-mediated knock-in cell-lines targeting the endogenous *Kat6b* and *Kat7* loci with a HaloTag® fusion for live-cell imaging (Figure 1G and Suppl. Figure 1I). We chose *Kat6b* over *Kat6a* because of its higher mRNA levels in mESCs (Suppl. Figure 1H). Live-cell imaging of the HaloTag fusions in dividing mESCs, revealed that both KAT6B and KAT7 proteins remained partially associated with chromatin throughout mitosis (Figure 1G), but to different degrees and with distinct patterns. In agreement with the mass-spec results, KAT6B showed a stronger association with mitotic chromatin compared to KAT7. More specifically, KAT6B displayed robust enrichment along condensed metaphase chromosomes, suggesting widespread mitotic binding or chromosome coating. In contrast, KAT7 exhibited a more restricted distribution at specific regions on mitotic chromosomes (Figure 1G-H).

Collectively, these data challenge the prevailing view that histone acetyltransferases are broadly excluded from mitotic chromatin. Instead, our findings demonstrate that KAT6A/B, KAT7, and multiple components of their catalytic complexes remain associated with mitotic chromosomes alongside their target acetylation marks. The coordinated retention of these enzymes and their catalytic products support a model in which histone acetylation is actively maintained during mitosis and contributes to the preservation of regulatory information across cell division.

### Site-specific retention of KAT6/7 binding and activity during mitosis

The different extents and patterns of KAT6B and KAT7 mitotic chromatin association observed in our live-cell imaging, suggest potentially distinct genomic occupancy. To address this, we performed chromatin immunoprecipitation (ChIP-seq) in both asynchronous and highly synchronized mitotic ESCs. This identified thousands of high-confidence KAT6B and/or KAT7 binding sites in asynchronous cells (KAT6B = 54,880, KAT7 = 11,844), with many KAT6B-specific peaks, but also many shared targets (9661) that were predominantly located around promoters (Suppl. Figure 2A-C). Intriguingly, metaplot analysis centered on the TSS of their common target genes revealed slightly different occupancy patterns: while KAT7 was mostly bound at the +1 nucleosome and depleted from the nucleosome-free promoter region, KAT6B showed broader binding around the TSS (Suppl. Figure 2D). These observations reveal that KAT6B and KAT7 have both overlapping and distinct genomic targets, suggesting both redundant and unique roles in regulating acetylation and target gene expression.

We then focused on mitotic ESCs, which showed a global reduction in both KAT6B and KAT7 binding, with 10,427 detected peaks for KAT6B (19% of asynchronous peaks) and only 689 peaks for KAT7 (5% of asynchronous peaks) (Suppl. Figure 2E-G), consistent with the different degrees of mitotic retention observed in our live-cell imaging and mass-spec experiments. By plotting the relative change of KAT6B ChIP-seq signal (log2FC(Mitotic/asynchronous signal)) versus the asynchronous signal (either normalized by sequencing depth or by spike-in), we could classify peaks into three different clusters based on their degree of mitotic retention: Lost (log2FC < -1.9), Low-retention (log2FC > -1.9 and < -0.8) and High-retention (log2FC >-0.8) (Figure 2A, Suppl. Figure 2G-H and Suppl. Table 3). Importantly, most of the strongest asynchronous peaks fell into the “Lost” peak category, arguing against a simple proportional persistence of the signal and instead for a regulated, locus-specific mechanism of mitotic retention (Suppl. Figure 2I). By projecting the KAT7 asynchronous and mitotic ChIP-seq signals onto the KAT6B peak categories, we observed some degree of KAT7 mitotic binding, especially around the KAT6B High-retention peaks (Figure 2B), suggesting potential cooperativity in their mitotic retention at these regions. Parallel profiling of the target histone marks of KAT6B and KAT7, H3K23ac and H3K14ac, in asynchronous and mitotic cells showed strong mitotic co-occupancy with both the High- and Low-retention KAT6B peak clusters, and substantially weaker signal around the lost KAT6B peaks (Figure 2C). These results suggest that persistent mitotic binding of KAT6B and, secondarily, of KAT7 might be responsible for the mitotic retention of these histone acetylation marks.

**Figure 2:**
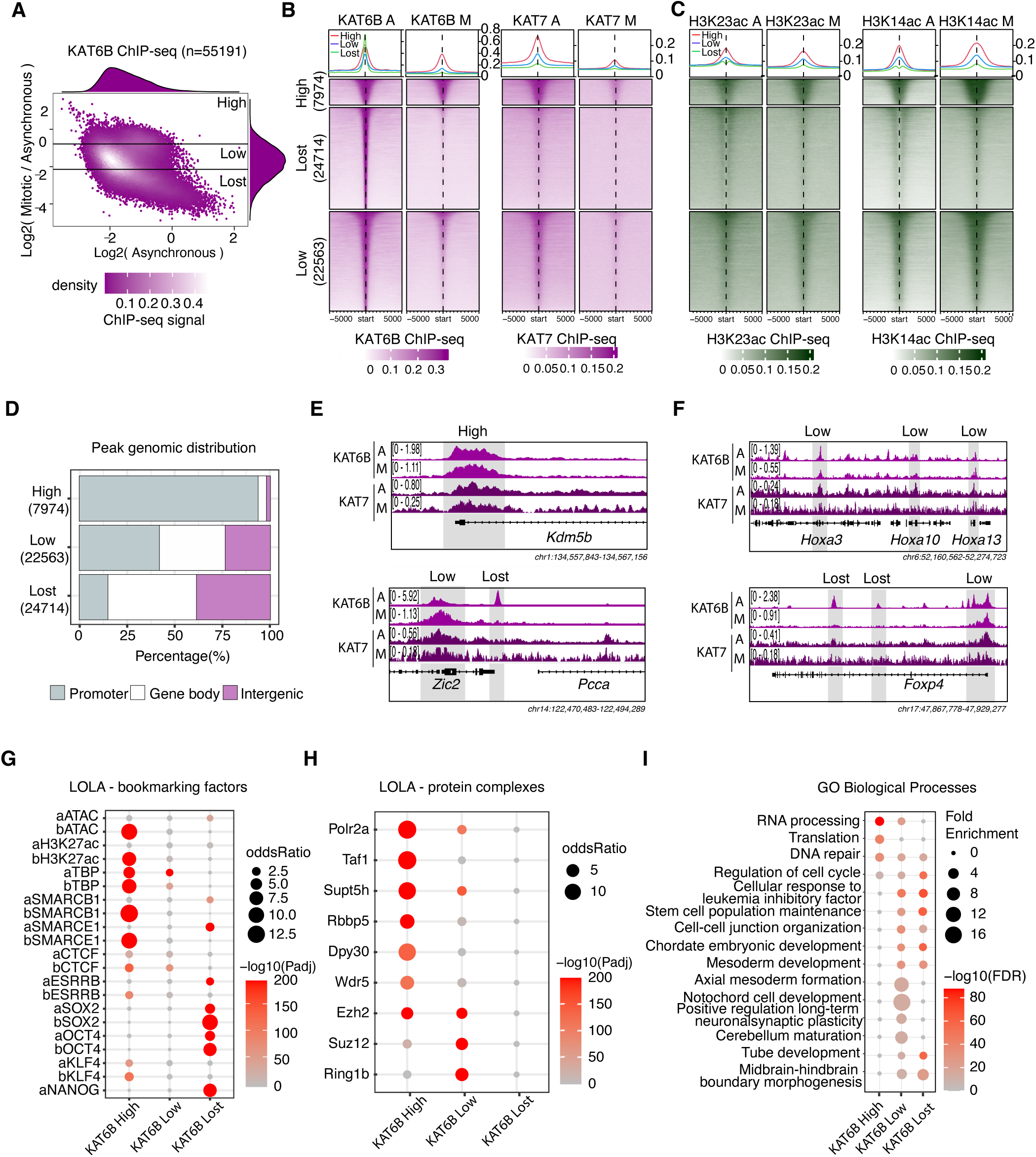
KAT6B, and to a lesser extent KAT7, and their target acetylation marks remain bound at developmental promoters during mitosis. **A.** Scatterplot comparing the relative change in KAT6B binding during mitosis (expressed as log2 fold change of mitotic versus asynchronous ChIP-seq signal after normalization over input and sequencing depth) over the asynchronous signal. The histograms along the axes of the plot depict the density of ChIP-seq ratios and signal and were used as a reference to set the cut-offs for different retention levels: high (log2FC > -0.8), low (log2FC > -1.9 and < - 0.8) retained peaks and lost (log2FC < -1.9) peaks. (See also similar plot in Suppl. Fig 2g with spike-in normalization and Suppl. Table 3). **B.** Tornado plots showing the asynchronous and mitotic KAT6B and KAT7 ChIP-seq signals (normalized over input and sequencing depth) for the different peak categories based on the relative KAT6B retention levels, as defined in (a) (high = red, low = blue, lost = green). The average signal of n= 2 independent biological replicates is projected +/- 5 kb around peak centers. **C.** Same as (b) but for asynchronous and mitotic H3K14ac and H3K23ac ChIP-seq signals (normalized over input and sequencing depth). The average signal of n= 2 independent biological replicates is projected +/- 5 kb around peak centers. Of note, the asynchronous H3K14ac ChIP-seq signal is based on published datasets^82^. **D.** Stacked bar plot representing the genomic distribution (percentages) of the different KAT6B ChIP-seq peak categories as defined in panel (a) across promoters, gene bodies or intergenic regions. **E-F.** Representative genome browser snapshots of KAT6B and KAT7 binding on asynchronous and mitotic cells at different genomic regions. Examples display high-retention and low-retention peaks, at the *Kdm5b* and *Zic2* promoters, respectively, the low-retention developmentally-relevant *Hoxa* cluster, and the *Foxp4* locus, with a low-retaintion peak at the promoter many lost intragenic peaks. IGV tracks are shown auto-scaled The relative ChIP-seq intensities are shown on the left of each track, while the exact genomic coordinates are shown at the bottom of each example. **G.** Dot plot showing the relative enrichment of high-, low-retention and lost KAT6B peaks (as defined in panel **(a)** for previously described bookmarking factors using published asynchronous/interphase and mitotic peaks datasets^83^. Prefix ‘a’ stands for asynchronous-specific peaks and ‘b’ stands for bookmarked peaks. **H.** Dot plot showing top enriched proteins at the high-, low-retention and lost category of peaks based on unbiased ChIP enrichment analysis using the LOLA tool and a curated set of published ChIP-seq for mouse ESCs. Proteins belonging to the Pol2, MLL and PRC complexes are shown as examples. The full LOLA analysis is provided in Supplementary Table 4. **I.** Dot plot showing top enriched Gene Ontology biological processes for each KAT6B peak category using the GREAT tool with basal plus proximal extension (5 kb upstream, 1 kb downstream) and distal extension up to 250 kb. The full list of gene ontology terms is shown in Table 4.

We then checked the genomic distribution of the different KAT6B peak clusters. We observed that the High-retention cluster contained almost exclusively promoter regions (∼90%) in sharp contrast with the Lost peaks that were predominantly distal to promoters (within gene bodies or intergenic regions) (Figure 2D), consistent with prior studies which report stronger mitotic retention on promoters^12,56^. However, the Low-retention peaks were located both around promoters (∼40%) and at distal genomic regions (Figure 2D). Genome Browser examples for each peak category are shown in Figure 2E-F. ChIP-enrichment analysis for each KAT6B peak cluster using published ChIP-seq datasets from mouse ESCs revealed a significant and preferential association of the KAT6B High-retention peaks with previous mitotic bookmarks, including mitotically retained ESRRB, TBP, SMARCB1 and SMARCE1^54,57–59^ as well as bookmarked ATAC-seq and H3K27ac regions^12,32^ (Figure 2G and Suppl. Table 4). Unbiased ChIP-enrichment analysis also showed that High-retention KAT6B peaks were strongly enriched for binding of RNA Pol2 and MLL-complex subunits, while regions with Low-retention were significantly associated with components of the Polycomb Repressive Complexes (PRC1/2) (Figure 2h. i.e., EZH2, SUZ12, RING1B). This suggests that KAT6B bookmarks both active and bivalent/poised promoters during mitosis. In agreement, Gene Ontology (GO) analysis showed that although highly bookmarked peaks were significantly enriched in chromatin organization and gene expression categories, the low-retention group was strongly associated with genes involved in stem cell population maintenance and various developmental processes (Figure 2I and Suppl. Table 4). Interestingly, Lost peaks were also enriched for both stem cell and differentiation categories (Figure 2I).

Together, these data and analyses document the selective mitotic retention of KAT6B, and to a lesser degree of KAT7, along with their histone acetylation targets, suggesting potential bookmarking activity. Importantly, in addition to the strong mitotic retention around active promoters, we also detected a weaker mitotic binding on poised enhancers and promoters, suggesting potential roles both in the transmission of an “active” transcriptional memory but also in the priming of developmental programs.

### Dual depletion of KAT7 and KAT6B results in H3K14ac loss and partial eviction of bookmarking factors from mitotic chromatin without affecting accessibility

The co-occurrence of mitotic KAT6B and KAT7 binding with different histone acetylation marks and other bookmarking proteins suggests a potentially active role for these HATs for maintaining mitotic chromatin acetylation and composition. To directly test this, we engineered individual and dual KAT6B and KAT7 degron mESC lines, by CRISPR-mediated knock-in (Figure 3A). Specifically, we introduced in-frame with the C-termini of the endogenous *Kat6b* and/or *Kat7* loci a cassette that encodes for the FKBP12 peptide^60^ (for dTAG-induced degradation) and HA-tags (for detection and pull-down with anti-HA antibodies), and a transcriptional reporter (P2A-mCherry or P2A-EGFP, for prospective isolation of targeted cells) (Figure 3A and Suppl. Figure 3A). After genotyping (Supp. Figure 3B), we confirmed the complete and rapid (by 1hr) dTAG-induced depletion of tagged proteins in different, clonal homozygous lines both by Western Blot (Figure 3B and Suppl. Figure 3C) and intracellular FACS (Suppl. Figure 3D).

**Figure 3.**
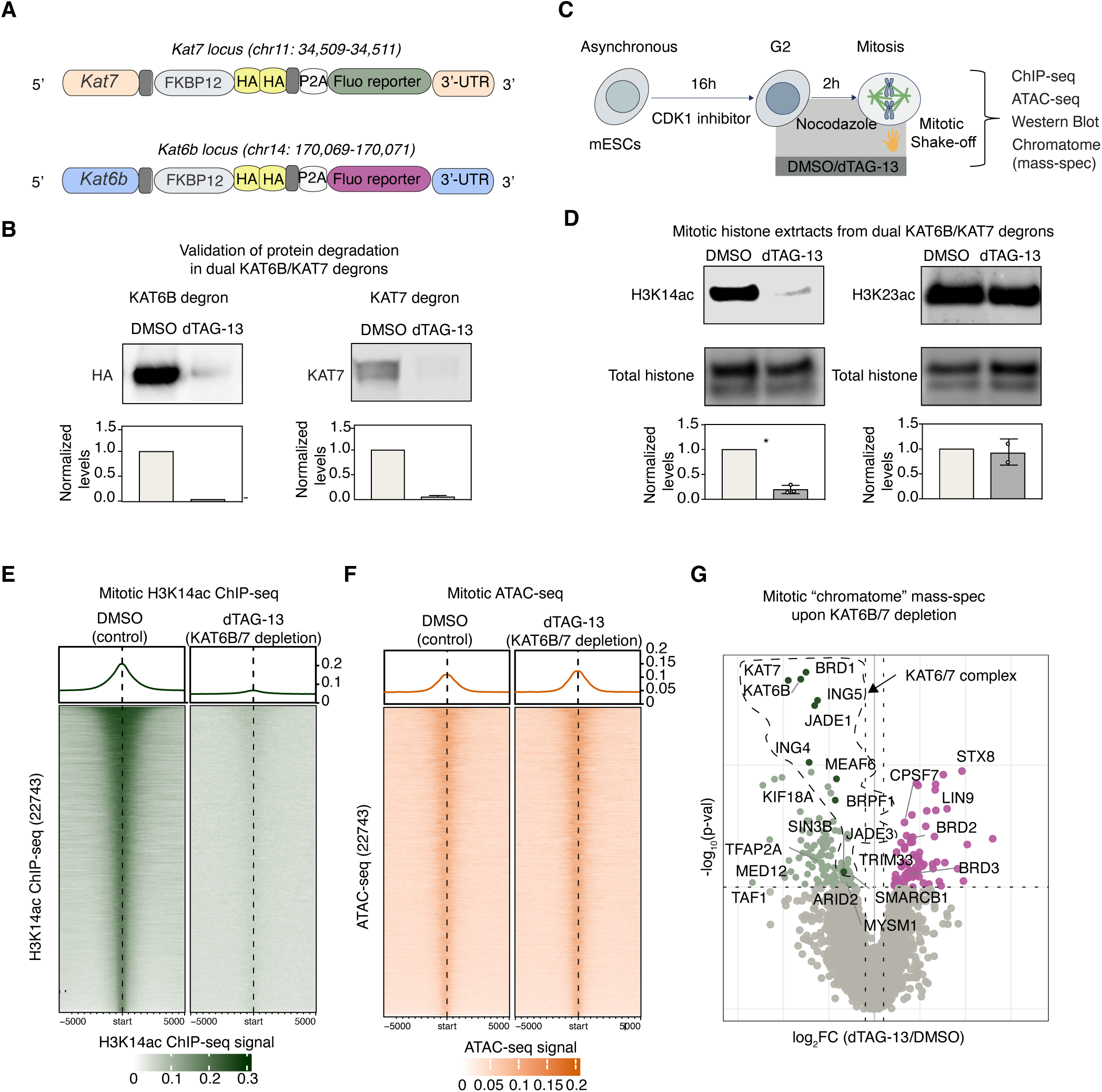
Dual depletion of KAT7 and KAT6B results in H3K14ac loss and partial eviction of bookmarking factors from mitotic chromatin without affecting accessibility. **A.** Schematic showing the knock-in cassettes used to generate individual and dual mouse KAT7 and KAT6B degrons. The knock-in cassettes were introduced into the endogenous *Kat7* and *Kat6b* gene loci in mESCs after their last exons. mKAT7 and mKAT6B proteins are fused to the FKBP12^F36V^ peptide for dTAG-13 mediated degradation, surrounded by linkers and 2xHA tags, while they were also transcriptionally linked through P2A to fluorescent reporter genes to facilitate cell-line generation. **B.** Western blot analysis showing nearly full depletion of KAT6B and KAT7 proteins upon dTAG-13 treatment in asynchronous cells. The antibody signal was normalized to a total protein stain. (not shown). **C.** Experimental strategy to characterize mitotic cells without KAT6B and KAT7 by ChIP-seq, ATAC-seq, Western Blot and chromatin proteomics. Dual-FKPB degrons were synchronized in mitosis by a dual-CDK1 inhibitor/nocodazole block and collected by mitotic shake-off. For acute depletion of the proteins specifically during M-phase, the DMSO/dTAG-3 treatment was added together with the nocodazole 2 hours prior to the shake-off collection. **D.** Western blot analysis of mitotic H3K14ac and H3K23 levels upon KAT6B/7 co-depletion in mESCs. Total histone staining is used as a loading control. Levels are normalized to the total histone and relative to the DMSO control. n= 2 biological replicates. H3K14ac levels were significantly lower after one-sample two-tailed t-test. *p < 0.05. H3K23ac levels did not show significant differences p > 0.05. **E.** Tornado plot representing mitotic H3K14ac ChIP-seq in DMSO and dTAG-13-treated cells in the dual-KAT6B/7 degron. 22743 peaks were called in the DMSO ChIP-seq. The average ChIP-seq signal of n = 2 technical replicates is shown after normalization over input and sequencing depth. **F.** Tornado plot representing the ATAC-seq signal projected on the 22743 peaks called in the panel (e) under DMSO or dTAG-13 treatment. Average depth-normalized signal from n = 2 technical replicates is shown. **G.** Volcano plot showing the relative enrichment or depletion of proteins on mitotic chromatin upon dTAG-13 treatment in the dual-KAT6B/7 degron relative to the DMSO controls as determined by quantitative mass spectrometry of chromatin-bound fractions. The log_2_fold change of (dTAG-13/DMSO) is represented on the x-axis and the -log_10_(p-val) in the y-axis. Proteins significantly depleted from mitotic chromatin are shown in green (N = 122; p val > 0.1, log2FC < 0) while proteins significantly enriched on mitotic chromatin are shown in magenta (N = 71; p-val > 0.1, log2FC > 0). KAT7 and KAT6B together with all the subunits forming their complexes are among the top depleted factors, and highlighted in darker green and surrounded by a dotted line. Other depleted proteins of interest are also highlighted in darker green. Differential analysis was performed using MAGMa^81^ on quadruplets (2 biological replicates (independent dual degron clones and 2 technical replicates from each clone). The full list of detected proteins is shown in Suppl. Table 2.

Next, we quantified the effects of 48h HAT depletion on their main target histone marks, H3K14ac and H3K23ac levels by Western Blot (Suppl. Figure 3E). KAT7 depletion alone led to a global reduction of H3K14ac levels by 70%, while the combination with KAT6B depletion resulted in a near-complete loss of this mark, with less than 10% of the WB signal remaining. Intriguingly, KAT6B depletion alone showed a slight but significant increase of H3K14ac (Suppl. Figure 3E), in agreement with recent observations in KAT6B knockout cells and mice^46,61^. These results support a complex interplay between these HATs in regulating H3K14ac. In contrast to the strong impact on H3K14ac, H3K23ac levels remained unaffected upon KAT7 and/or KAT6B degradation. This suggests that in our conditions other HATs, such as KAT6A, are primarily responsible for depositing this mark in mESCs or can compensate for the KAT7/KAT6B loss (Suppl. Figure 3E). Unbiased histone proteomics upon dual KAT7/6B depletion independently validated the strong and significant impact on H3K14ac (p-adj val < 0.1), with no other histone acetylation marks showing consistent changes (Suppl. Figure 3F).

Using our dTAG mESC lines, we then assessed the impact of dual KAT7/6B depletion specifically during mitosis. To do so, we employed our synchronization strategy and added DMSO/dTAG-13 treatment 2 hours prior to the mitotic shake-off (Figure 3C). Consistent with our results in asynchronous cells, we observed a strong reduction of the H3K14ac levels with no significant global changes of H3K23ac (Figure 3D). H3K14ac ChIP-seq upon mitotic KAT7/6B depletion (using human ESCs for spike-in normalization) confirmed the dramatic impact on H3K14ac with almost undetectable H3K14ac signal left genome-wide (Figure 3E and Suppl. Figure 3G), indicating that the two proteins are responsible for the active deposition of H3K14ac during mitosis. Of note, dual KAT7/6B degradation in asynchronous cells also induced a genome-wide reduction of H3K14ac but not a complete loss of the signal, suggesting some degree of compensation by other HATs in interphase cells but not in mitosis (Suppl. Figure 3H).

Considering the strong association of mitotic KAT6B/KAT7 binding with bookmarked ATAC-seq peaks (Figure 2G), we reasoned that KAT7/6B mitotic depletion could be accompanied by chromatin compaction. However, despite the near-complete loss of H3K14ac following KAT6B/KAT7 depletion, chromatin accessibility remained largely unchanged both genome-wide and at regions with acetylation loss (Figure 3F and Suppl. Figure 3G). These observations suggest that mitotic KAT6/7 binding and KAT6/7-dependent acetylation do not primarily function through maintaining chromatin accessibility.

Finally, we investigated whether acute depletion of KAT7 and KAT6B (2 hr) affected the protein composition of mitotic chromatin by quantitative mass-spectrometry. Compared with the DMSO control samples, dTAG treatment resulted in a significant reduction of 122 proteins from the mitotic chromatin and a relative enrichment of 71 proteins (Figure 3G). As expected, KAT6B and KAT7 were among the top depleted factors (Figure 3G). Notably, along with KAT6B and KAT7, all major subunits from their core MOZ/MORF and HBO1 complexes, respectively, were also significantly evicted from mitotic chromatin, indicating that KAT6B and KAT7 are critical for the mitotic chromatin composition and assembly of these complexes. Finally, acute KAT6B/KAT7 depletion destabilized not only the HAT complexes themselves but also reduced the association of additional chromatin regulators with mitotic chromatin, including the bookmarking factors TAF1^62^ and SMARCB1^54^ as well as the SWI/SNF component ARID2.

Together with the preserved chromatin accessibility observed upon KAT6B/KAT7 depletion, these findings support a model in which mitotic retention of KAT6/7 binding and/or H3K14ac contributes primarily to regulating/preserving the composition of the mitotic chromatome rather than simply maintaining an open chromatin state.

### Acute KAT7 and KAT6B co-depletion partially impairs post-mitotic transcriptional activation

The mitotic retention of KAT6B -and to a lesser extent of KAT7- and their role in maintaining and propagating H3K14ac during cell division could suggest a *bona fide* bookmarking function. If this is true, their depletion during M-to-G1 would cause dysregulation of post-mitotic transcriptional activation of bookmarked genes. To investigate this, we degraded KAT6B/KAT7 in mESCs undergoing mitotic arrest (as described above) and then released the cells into G1 for 3 hours in the continuous presence of DMSO or dTAG-13 (Figure 4A). After confirming that KAT7/KAT6B depletion had no effects on cell cycle progression and enabled efficient release in G1 (>83%) (Suppl. Figure 4A), we performed RNA-seq in G1 populations comparing DMSO and dTAG treatment conditions. This experiment detected hundreds of genes (n=678) that failed to be properly reactivated in the absence of KAT7 and KAT6B, showing significantly (p-adj val<0.05) lower levels under dTAG conditions, while only 78 genes were activated at higher levels (Figure 4B and Supp. Table 5). Importantly, over 90% of the downregulated genes overlapped with bookmarked KAT6B peaks (Low or High-retention), while these percentages were significantly lower among the unaffected (∼50%) or the upregulated genes (∼80%) (Figure 4C). Moreover, the downregulated gene set showed a significant overrepresentation of early activated genes during M-to-G1 transition, based on a previous time course study^12^, suggesting a selective impact on early activation (Suppl. Figure 4B). These results suggest that mitotic bookmarking by KAT6B/7 might actively promote the proper post-mitotic activation of these genes. In support of this notion, independent RT-qPCR analysis of pre-mRNAs for several of these DEGs showed significantly lower levels of nascent transcription as early as 1 hour after mitotic release in the absence of KAT6B/7, while non-bookmarked control genes showed proper reactivation (Figure 4D, non-bookmarked control: *Egfr*). Interestingly, GO analysis of the downregulated genes revealed a strong enrichment for processes related to stem-cell population maintenance, but also for various developmental processes, including morphogenesis, patterning, and cell fate specification, mirroring the gene ontology enrichment for the KAT6B/7-bookmarked genes (Figure 4E and Figure 2I).

**Figure 4.**
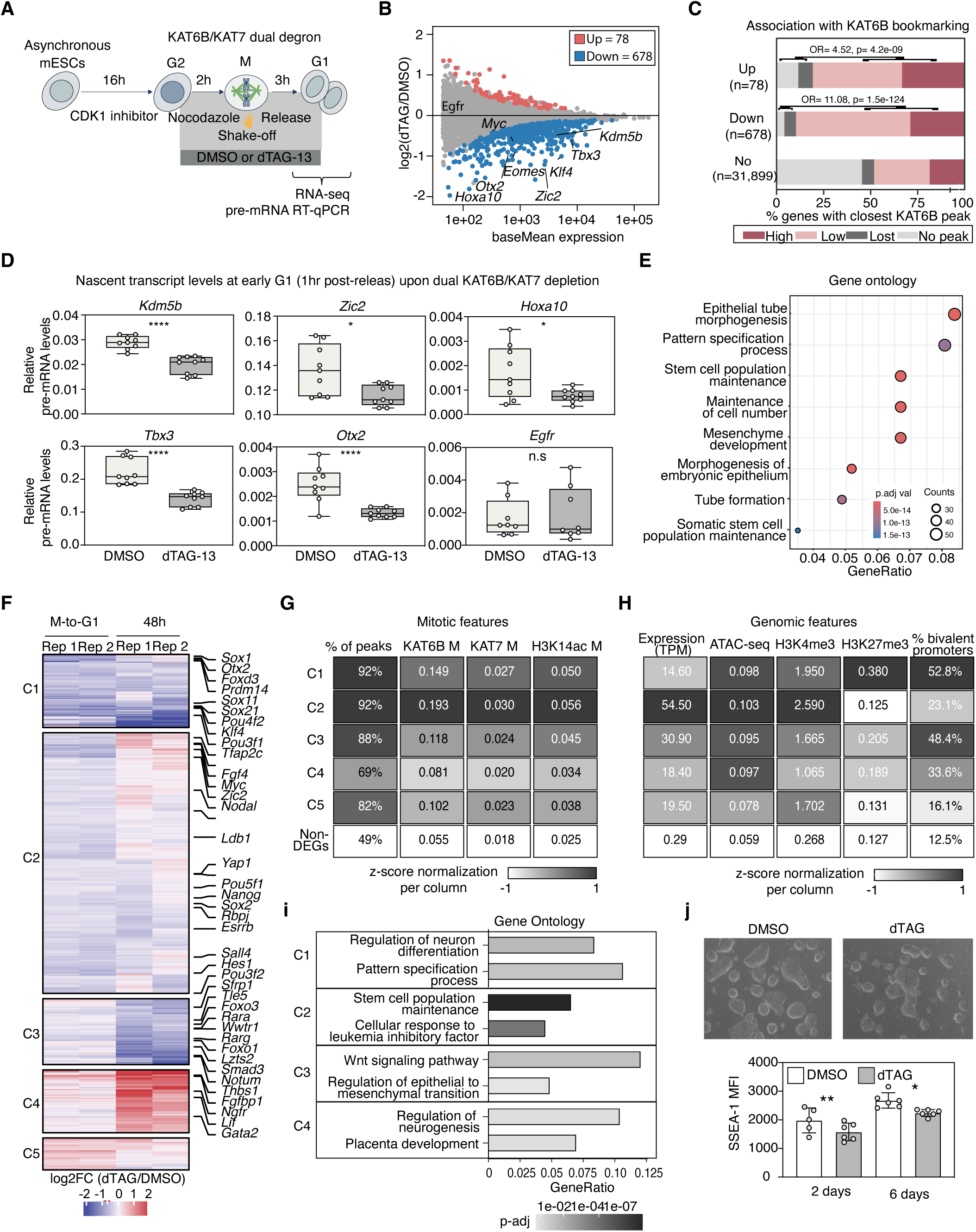
KAT6B and KAT7 co-depletion partially impairs post-mitotic transcriptional activation of stem-cell and lineage-specific genes. **A.** Experimental strategy to analyze transcriptional changes in G1 upon acute co-depletion of KAT6B/7. Mouse ESCs were synchronized in mitosis as previously described and released into G1 after mitotic shake-off for 1 and 3 hours. DMSO/dTAG-13 treatment was added 2 h prior to the mitotic shake-off and throughout the release into G1. G1 cells were subjected to RNA-seq and pre-mRNA RT-qPCRs. **B.** M/A plot of differentially expressed genes in DMSO versus KAT6B/7 co-depleted G1 cells (3 hours). The log_2_ fold change (dTAG13/DMSO) is plotted on the y-axis against the average expression levels for each gene (baseMean) on the x-axis for DMSO (n = 2 technical replicates). p-adj val > 0.05 was uses as a cut-off to call differential expression. Each point represents a single gene. Up-regulated genes are colored in red (N = 78) and downregulated genes are colored in blue (678). Non-DEGs are grey. Names of representative genes are depicted. The full list of DEGs is shown in Suppl.Table 5. **C.** Stacked bar plot showing the percentage of upregulated, downregulated and non-altered genes associated with the different KAT6B ChIP-seq categories, as defined in Fig. 2a. Overlap with high - and low-retention KAT6B peaks is shown in dark and pale pink, respectively, while overall with lost peaks or lack of any KAT6 peaks are shown in light and dark grey. The relative enrichment for bookmarked peaks between upregulated, downregulated genes and non-DEGs was calculated using a two-sided Fisher exact test, and the odds ratio and p-values are shown on top of each bar. **D.** Reverse-Transcriptase quantitative PCR analysis assessing the relative pre-mRNA levels (with qPCR primers overlapping intron-exon expansions) of select KAT6B-bookmarked (*Tbx3, Kdm5b, Zic2, Otx2 and Hoxa10*) or non-bookmarked (*Egfr*) genes after 1 h release into G1 in the presence of dTAG or DMSO. Box plots show expression levels for each gene relative to a housekeeping gene (*Hprt*) in DMSO and dTAG-13 (n = 3 biological replicates, N = 3 technical replicates). Statistical significance was assessed by two-sided unpaired t-test (* < 0.05, **** < 0.0001). Primers are provided in Suppl. Table 6. **E.** Gene ontology analysis showing top, select biological processes enriched for genes downregulated in G1 (as defined in panel b). The full list of gene ontology analysis results related to this figure is provided in Suppl. Table 4. **F.** Heatmap comparing the transcriptional changes upon KAT6/7 dual depletion either during M-to-G1 or in asynchronous mESC for 48 hr expressed as log2FC of RNA-seq of dTAG vs DMSO. Only genes that scored as significantly affected (DEGs) in either condition (see Fig. 4b or Suppl. Fig.4c) are plotted. Genes are clustered in 5 groups with K-means clustering (see full lists in Suppl. Table 6). Downregulated genes are color-coded in blue and upregulated genes are red. **G.** Bookmarking properties of each gene cluster depicted in panel (f) compared to non-DEGs. A heatmap per column is used to represent (i) % of KAT6B-bookmarked peaks per cluster (ii) Average depth-normalized mitotic signal for KAT6B, (iii) average depth-normalized mitotic signal for KAT7, (iv) average depth-normalized mitotic signal for H3K14ac. n = 2 biological replicates. Each column is normalized to their maximum and minimum values (z-score -1/+1). **H.** Genomics features defining each gene cluster depicted in panel (f) compared to non-DEGs based on in-house or published datasets from mouse ESCs^54,63^. A heatmap is used to represent per column (i) average expression levels of the genes in asynchronous mESCs (ii), average depth normalized ATAC-seq signal in asynchronous mESCs, (iii) average depth normalized H3K4me3 ChIP-seq signal in asynchronous cells (GSE189451), (iv) average depth normalized H3K27me3 ChIP-seq signal in asynchronous cells (GSE189561), (v) % of previously described bivalent promoters within the clusters^63^. Each column of information is normalized to their maximum and minimum (z-score -1/+1). **I.** Gene ontology analysis showing top two biological processes enriched per gene cluster as defined in panel (f). Terms are shown in a bar plot with the x-axis representing the GeneRatio and color-coded in different shades of grey based on p-adj value. Cluster 5 did not yield any significant terms. The full list of gene ontology analysis results related to this figure is provided in Suppl. Table 4. **J.** Top: representative images of mESCs colonies in control and dTAG-treated cells after 6 days of treatment. Bottom: Bar plot analysis showing the percentage of mESCs expressing the pluripotency surface marker SSEA-1 based on flow cytometry analysis (n = 2 biological clones, N = 3 technical replicates per clone) after two or six days of treatment with dTAG-13 compared to DMSO control cells. Asterisks indicate statistical significance as calculated by paired two-sided t-test (* p < 0.05, ** p < 0.01).

### Prolonged KAT7 and KAT6B co-depletion results in partial dysregulation of stem-cell-related genes and silencing of primed lineage-specific programs

To determine whether the transcriptional defects induced by acute KAT6B/7 depletion during the M-to-G1 transition persist after prolonged depletion, we performed RNA-seq following 48 h of dTAG treatment in asynchronous mESCs. Surprisingly, despite the robust transcriptional response observed during M-to-G1 (n=756 DEGs, see Figure 4B), prolonged depletion resulted in substantially fewer transcriptional changes (n=262 DEGs), including 145 downregulated and 117 upregulated genes (Suppl. Fig. 4C and Suppl. Table 5). This suggested that different classes of KAT6B/7 target genes exhibit distinct temporal responses, potentially reflecting compensatory mechanisms during interphase.

To identify distinct temporal patterns, we integrated the M-to-G1 and 48 h datasets and performed K-means clustering of all significant DEGs based on their transcriptional responses across both conditions (Figure 4F, Suppl. Figure 4E and Suppl. Table 6). Five major clusters emerged. The three largest clusters consisted predominantly of genes initially downregulated during M-to-G1. While Cluster 1 and Cluster 3 genes were further downregulated after prolonged depletion, Cluster 2 genes showed substantial recovery despite its initial sensitivity to KAT6B/7 loss. In contrast, Clusters 4 and 5 contained genes that were transiently or persistently upregulated.

To investigate the molecular basis underlying these distinct transcriptional behaviors, we compared the chromatin features associated with each cluster. Consistent with our earlier analyses, 88–93% of genes within the three downregulated clusters (Clusters 1–3) were associated with KAT6B-bookmarked promoters and displayed significantly stronger mitotic occupancy of KAT6B, KAT7, and H3K14ac than either upregulated genes (Cluster 4) or non-differentially expressed genes (Figure 4G). Interestingly, Cluster 5 genes, which were transiently upregulated following acute KAT6B/7 depletion, also showed strong KAT6B bookmarking (82%) but lower mitotic levels of H3K14ac, suggesting that the coexistence of KAT6B/7 occupancy and high H3K14ac levels dictate the direction of transcriptional regulation. We next examined promoter chromatin states and basal transcriptional activity to identify features distinguishing the downregulated clusters. Whereas Cluster 2 genes exhibited the hallmarks of active promoters, including high accessibility and H3K4me3 levels, and robust transcription, Cluster 1 genes were characterized by the concurrent enrichment of H3K4me3, H3K27me3, and persistent H3K14ac, consistent with a poised chromatin configuration. Cluster 3 displayed an intermediate profile, with lower levels of both H3K4me3 and H3K27me3. Accordingly, Clusters 1 and 3 were expressed at significantly lower levels than Cluster 2 and were enriched for previously annotated bivalent developmental genes^63,64^, including *Hoxa10, Nkx6-2, Gata3,* and *Otx2*. The coexistence of H3K14ac with H3K4me3 and H3K27me3 raises the possibility that these loci adopt a trivalent chromatin state -as reported previously in asynchronous cells^65^-that is maintained through mitosis in a KAT6B/7-dependent manner. In agreement with their chromatin features and transcriptional levels, Gene ontology analysis revealed that Cluster 2 was highly enriched for stem-cell maintenance and pluripotency regulators, including *Sox2, Nanog,* and *Pou5f1*, whereas Cluster 1 preferentially contained genes involved in developmental patterning, cell-fate specification, and neuronal differentiation (Figure 4I and Suppl. Table 6).

Together, these findings identify two fundamentally different classes of KAT6B/7 target genes. Highly expressed pluripotency genes are acutely sensitive to KAT6B/7 loss during mitotic exit but largely recover following prolonged depletion, suggesting that their regulation can be buffered by compensatory mechanisms during interphase. Consistent with this interpretation, prolonged KAT6B/7 depletion caused only a modest but significant reduction of the pluripotency marker SSEA1 after two and six days of dTAG treatment (Figure 4J), indicating that the pluripotency program is largely preserved despite its transient disruption during mitotic exit. In contrast, KAT6B/7 bookmarking and activity is very critical for lowly expressed developmental genes, which exhibit progressively stronger transcriptional defects upon prolonged depletion. These observations support that KAT6B/7 contribute both to the faithful post-mitotic resetting of the self-renewal program and the maintenance of developmental competence by priming lineage-associated genes for future activation.

### KAT6/7 mitotic bookmarking and roles in post-mitotic transcriptional resetting are conserved in human ESCs

Given the established relevance of KAT6/7 factors to neuronal differentiation and human neurodevelopmental disorders^42,44–50^, we sought to determine whether KAT-dependent mitotic bookmarking is conserved in human embryonic stem cells (hESCs) and contributes to neuronal lineage differentiation, leveraging robust and well-characterized directed differentiation protocols. First, using mass spectrometry in asynchronous and mitotic hESCs (Figure 5A), we validated both the broad mitotic retention of histone acetylation (Suppl. Figure 5A-D and Suppl. Table 1) and of KAT6/7 proteins and various components of the MOZ/MORF and HBO1 complexes (Suppl. Figure 5E and Suppl. Table 2). As an independent validation, live imaging of the endogenous Halo-tagged KAT6B and KAT7 proteins in dividing hESCs revealed a persistent association of both proteins with mitotic chromosomes (Figure 5B and Suppl. Figure 5F-H). These results support a conserved role for KAT6/7-containing HAT complexes in transmitting active epigenetic memory in undifferentiated pluripotent stem cells.

**Figure 5.**
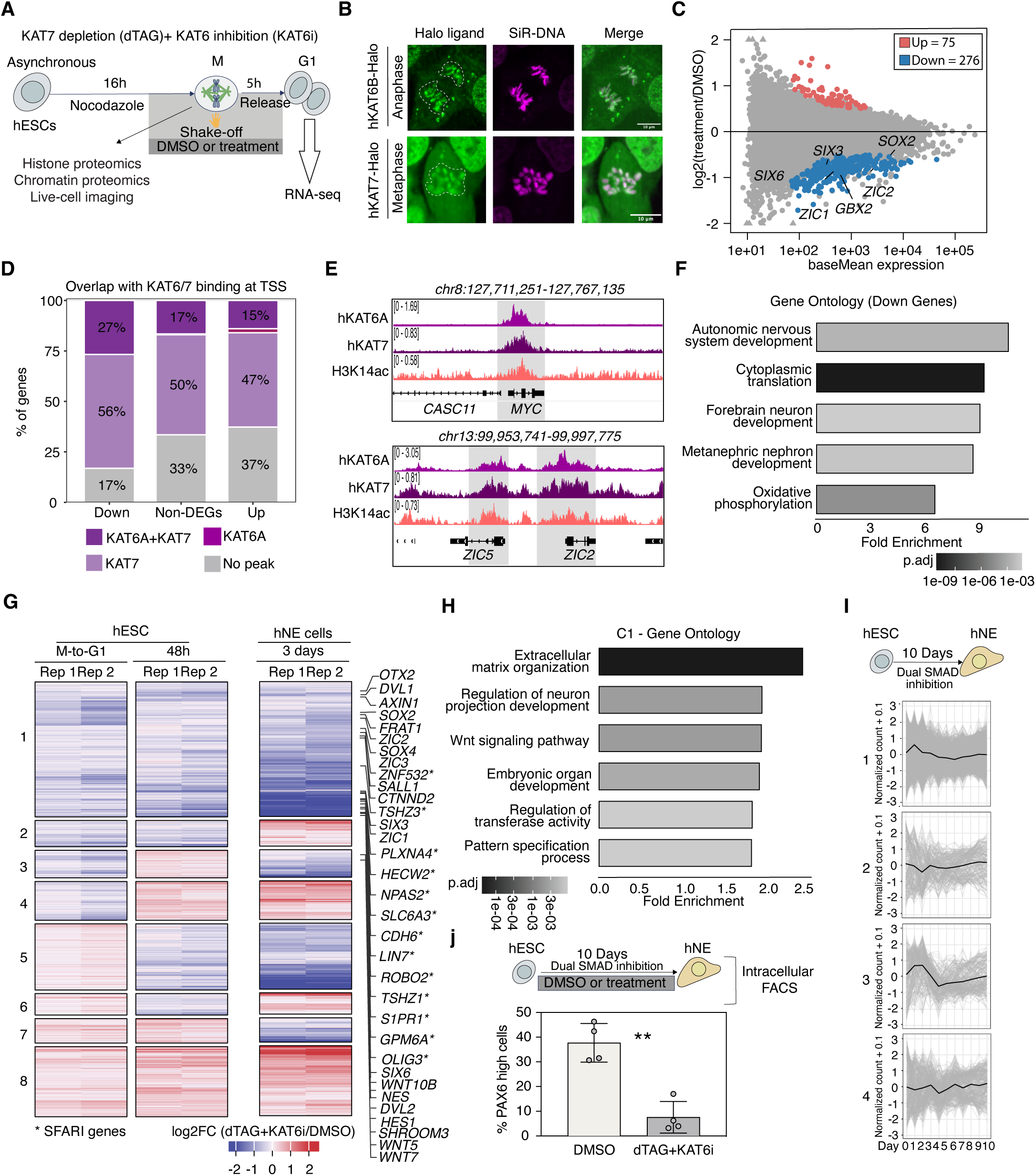
KAT6/7 mitotic bookmarking and roles in post-mitotic transcriptional resetting are conserved in hESCs. **A.** Experimental strategy for the synchronization and isolation of mitotic human ESCs (hESC) for histone and chromatin proteomics analysis by mass-spectrometry in comparison with asynchronous cells and to assess post-mitotic transcriptional resetting in human ESCs upon KAT7 depletion and KAT6 inhibition. Human ESCs were synchronized in mitosis by a 16 h nocodazole treatment, collected by mitotic shake-off and either subjected to histone and chromatin proteomics or release into G1 for RNA-seq. **B.** Live-cell imaging of hESCs HaloTagged for KAT6B and KAT7. Representative images show the HaloTagged protein in green in different mitotic phases. DNA is shown in magenta. Scale bar = 10 μm. **C.** M/A plot of differentially expressed genes in dTAG+KAT6i-treated cells vs DMSO control cells in G1. The log_2_ fold change (treated/DMSO) is plotted on the y-axis against the average expression for each gene (baseMean) on the x-axis for DMSO samples (n = 3 technical replicates). p-adj val > 0.05 was used as a cut-off to call differential expression. Each point represents a single gene. Upregulated genes are colored in red (N = 75) and downregulated genes are colored in blue (276). Non-DEGs are grey. Names of representative genes are depicted. The full list of DEGs is shown in Suppl. Table 5. **D.** Stacked bar plot showing the percentage of DEGs and non-DEGs with promoter peaks for either KAT6A and KAT7, KAT7 alone or KAT6A alone. Downregulated genes showed a significant overrepresentation of KAT6A + KAT7 common peaks (p = 2.73e-05) and underrepresentation of no-associated KAT6/7 peaks (p = 1.57e-09) compared to non-DEGs, as assessed by a two-sided Fisher exact test. **E.** Genome browser tracks showing how the promoters of representative downregulated DEGs (as defined in (b)) overlap with binding of KAT6A (light purple), KAT7 (purple) and H3K14ac (salmon pink), as assessed by ChIP-seq in hESCs. The depth-normalized ChIP-seq signals are shown on the left of each track, while the genomic cordiates of each losus are shown on the top. **F.** Bar plot showing top enriched gene ontology categories in the G1 downregulated genes. The full list of terms is provided in Suppl. Table 4. **G.** Heatmap comparing the transcriptional changes upon KAT6/7 dual perturbation (dTAG+KAT6i) either during M-to-G1 or in asynchronous hESC for 48 h or at day 3 of neuroectoderm specification, expressed as log2FC of RNA-seq of (dTAG+KAT6i) vs DMSO. Only genes that scored as significantly affected (DEGs) in either condition (see Fig. 6b or Supp. Figure 6i) are plotted. Genes are clustered manually into 8 clusters based on their log2FC trends across conditions. Downregulated genes are color-coded in blue while upregulated genes are color-coded in red. Some representative genes are shown for clusters 1 and 3. Stars mark genes related to autism-spectrum disorders as reported in the SFARI database^69^. **H.** Bar plot showing the top enriched gene ontology biological processes in Cluster 1 of the heatmap shown in (g). x-axis show the geneRatio per category and y-axis the GO terms. P-adj value for each category is shown in a grey gradient. The full list of terms per cluster is provided in Suppl. Table 4. **I.** Normalized scRNA-seq counts for gene clusters 1-4 as defined in panel (g) showing their expression kinetics during a previously published hESC-to-neuroectoderm differentiation time course (Xu et al, 2025) (see full gene lists in Suppl. Table 6). Each panel represents an individual gene cluster. Gray lines indicate normalized expression trajectories for each individual gene while the black line represents the cluster mean. Cluster 1 and 3 specifically show an early upregulation at day 1 or 2, suggestive of genes that need to be upregulated or activated in the initial stages of differentiation. **J.** Analysis of the effects of KAT6/7 perturbation on neuroectoderm specification. *Top:* experimental strategy to differentiate hESCs into neuroectoderm within 10 days by dual Smad inhibition. Over the course of differentiation, starting at Day 0, the cells were either kept in DMSO or in media containing dTAG-v1 to deplete KAT7 and 100 nM of KAT6 inhibitor. At day 10, the levels of PAX6, a neuroectoderm marker, was anaylzed by intracellular FACS. *Bottom:* bar plot depicting the % of PAX6-high expressing cells after 10 days of neuroectoderm specification under KAT6/7 perturbation. n= 2 biological replicates and N = 2 technical replicates were quantified by intracellular FACS for each condition. Under treatment, the % of PAX6-high cells is significantly lower compared to DMSO cells as assessed by a paired two-sided t-test. ** p < 0.01.

Next, we investigated the transcriptional consequences of their disruption during M-to-G1. To do so, we generated and validated a KAT7 degron hESC line (Suppl. Figure 6A) and combined it with a commercially available catalytic inhibitor (KAT6i)^66^ targeting both, KAT6A and KAT6B. These cells underwent mitotic arrest-and-release into G1 in the continuous presence of dTAGv1+KAT6i (Figure 5A) or DMSO as a control, followed by collection after 5 hours of release (as hESCs exhibit a longer G1 compared to mESCs). After we validated the efficiency of each treatment (i.e., KAT7 degradation and KAT6 catalytic inhibition by WB) and the successful release into G1 (by FACS analysis) (Suppl. Figure 6C-D), we performed RNA-seq to determine potential defects in the M-to-G1 transcriptional resetting. Consistent with our hypothesis, cells with KAT6/7 perturbation showed global transcriptional downregulation, indicating impaired reactivation of their transcriptional program. Specifically, we detected 276 downregulated genes (adjusted p-value< 0.05) while only 75 genes were upregulated (Figure 5C). Interestingly, we noticed that some of the affected genes overlapped with those identified in mESCs (i.e: *Zic2, Otx2, Sox2*) (Figure 4B), suggesting conserved regulatory targets for these HATs during M-to-G1. ChIP-seq analysis of KAT7 and KAT6A in hESCs revealed significantly more peaks and stronger binding of both proteins to the promoters of downregulated genes compared with upregulated or unaffected genes (Figure 5D-E and Suppl. Figure 6H), suggesting direct effects. Intriguingly, the unique binding footprints we detected for KAT7 and KAT6B around the TSS of their shared target genes in mouse ESCs were also observed in human KAT7 and KAT6A (Suppl. Figure 6F-G), further supporting both overlapping and distinct functions of these HATs.

Consistent with a role of KAT6/7 in regulating proper inheritance and reactivation of the stem cell program, KAT6/7 perturbation during M-to-G1 resulted in reduced transcriptional levels of various core pluripotency factors, including *NANOG, KLF4, MYC, POU5F1* and *SOX2* (Suppl. Figure 7A). In addition, GO analysis of the downregulated genes revealed a strong enrichment for developmental processes, such as pattern specification and autonomic nervous system development (Figure 5F). Notably, many of these genes are activated early during neuroectoderm differentiation, suggesting that KAT6/7 activity primes transcriptional programs required for subsequent neuronal lineage progression.

### KAT6/7-activity primes developmental signaling networks for neuronal lineage specification

To test the impact of KAT6/7 perturbation on both the maintenance of the stem cell program and the establishment of the neuroectoderm lineage, we subjected the KAT7-dTAG hESCs to prolonged dTAGv1 and KAT6i treatment either in self-renewal conditions (hESC 48h) or during neuroectoderm differentiation upon dual SMAD inhibition^67^ (NE d3). RNA-seq analysis identified thousands of differentially expressed genes (DEGs) in each condition relative to their corresponding DMSO controls (Suppl. Figure 7B), with downregulated genes outnumbering upregulated genes, consistent with the predominantly activating functions of KAT6 and KAT7.

To determine whether the acute transcriptional changes induced by KAT6/7 perturbation during the M-to-G1 transition (Figure 6B) contribute to the transcriptional defects observed upon prolonged inhibition, we integrated all significant DEGs (adjusted p-value < 0.05, G1 = 351, ES = 1596, NE = 3141) from the M-to-G1, hESC 48 h, and NE d3 datasets and clustered them according to their transcriptional responses across conditions (Figure 5G, Suppl. Figure 7C and Suppl. Table 6). Focusing on genes that were downregulated during the M-to-G1 transition, the largest cluster (Cluster 1; n = 1,035) displayed persistent downregulation upon prolonged treatment in self-renewing hESCs and even stronger suppression during neuroectoderm differentiation. This progressive phenotype suggests that KAT6/7 activity is required both to prime these genes in pluripotent cells and to support their full activation during lineage commitment. Another interesting cluster, Cluster 3, contained genes (n=211) that were affected transiently during M-to-G1, not at 48 h in hESCs, but then downregulated at day 3 of NE specification upon KAT6/7 perturbation, suggesting context-specific roles of these proteins. Importantly, Cluster 1 strongly enriched for signaling pathways and processes involved in morphogenesis, patterning and neuronal differentiation, including canonical Wnt signaling (i.e., *DVL1, AXIN1/2, FRAT1/2* etc) and neuroectoderm regulators (i.e., *SOX2, SOX4, SOX10, SALL1, HHEX, FOXD1* etc) (Figure 5H). On the other hand, Cluster 3 genes were mostly associated with oxidative phosphorylation and morphogenesis, including also WNT and polarity genes, such as *DVL2* and *SHROOM3* (Suppl. Table 6).

To independently assess the developmental relevance of these gene sets, we examined their expression dynamics during a previously published 10-day hESC differentiation time course toward neural progenitor cells (NPCs)^68^. Notably, genes belonging to Clusters 1 and 3 showed a clear trend for early induction during neuronal differentiation, whereas genes from other clusters displayed comparatively flat or heterogeneous expression patterns (Figure 5I). These observations further support the notion that KAT6/7 regulate transcriptional programs that are activated early during neuronal lineage commitment. Finally, both Clusters 1 and 3 included several genes with well-established roles in neurodevelopmental disorders, including 13 genes associated with autism spectrum disorders (e.g *LIN7A, CTNND2* and *OLIG3)*^69^, suggesting that the early function of KAT6/7 in regulating these genes could contribute to disease phenotypes.

Together, these findings support a model in which KAT6 and KAT7 activity during M-to-G primes developmental signaling, morphogenetic, and metabolic networks required for successful neuronal differentiation. Consistent with this model, continuous KAT6/7 perturbation (TAGv1 + KAT6i) throughout a 10-day hESC-to-neural progenitor cell (NPC) differentiation protocol resulted in a significant reduction in the fraction of PAX6-positive cells (Figure 5J and Suppl. Fig.7D), indicating impaired neuroectoderm specification. These results suggest that dysregulation of KAT6/7-dependent gene regulatory networks during M-to-G1 might translate into later neuronal defects and provide a potential mechanistic link between KAT dysfunction and neurodevelopmental disease.

## DISCUSSION

Mitotic bookmarking mechanisms are thought to preserve regulatory information across cell division, allowing daughter cells to rapidly re-establish transcriptional programs and maintain cellular identity^70^. While most prior work has focused on the retention of transcription factors and histone methylation modifications, especially of repressive H3K27me3 and H3K9me3 marks^32,71–73^, the contribution of histone acetylation to epigenetic memory has remained controversial^8^. The prevailing view is that, along with global transcriptional silencing, histone acetylation is drastically reduced during mitosis due to rapid turnover and HDAC activity. However, recent quantitative proteomic studies have challenged this view by demonstrating widespread retention of histone acetylation marks across mitosis in multiple cell types^30–32^. Our findings are fully consistent with these observations and extend them to both mouse and human pluripotent stem cells, where faithful transmission of cellular identity and developmental potential is particularly critical. Importantly, along with persistence of histone acetylation marks, our study also reports the mitotic retention and activity of various HATs, providing mechanistic evidence for the active maintenance of acetylation during cell division. Specifically, we characterize the mitotic retention of KAT6 family proteins and, to a lesser extent, of KAT7 and demonstrate that acute depletion of these enzymes results in near-complete loss of H3K14ac during mitosis. This is accompanied by selective disruption of chromatin-associated regulatory complexes and bookmarking factors, but remarkably little change in global chromatin accessibility. Together, these findings support a model in which HATs and histone acetylation function as integral components of mitotic chromatin, actively contributing to the preservation of regulatory information and chromatin composition across cell division.

One unexpected finding of our study is that mitotic acetylation may extend beyond simply marking active genomic regions. While mitotic bookmarking has traditionally been viewed through the lens of preserving ongoing transcriptional programs^14,70^, our data suggest an additional role in priming genes for activation during lineage commitment. Indeed, we observed partial mitotic retention of both KAT6 and H3K14ac on several bivalent, developmental genes, suggesting a link between mitotic acetylation and developmental competence. Gene networks sensitive to acute KAT6/7 perturbations were enriched for developmental signaling pathways, morphogenetic regulators, metabolic programs, and key components of canonical WNT signaling, many of which showed only modest transcriptional defects in pluripotent cells but progressively stronger perturbations during neuronal differentiation. Although KAT6 proteins have been reported to recognize and regulate bivalent developmental loci^74^ and neuronal programs^44–46^, our study documents their persistent binding and activity during mitosis and the M-to-G1 transition, an important window for cell fate specification. Here, we propose that the KAT6/7-dependent chromatin states established and maintained in self-renewing pluripotent cells facilitate the future activation of lineage-specific genes, thereby coupling mitotic chromatin inheritance to developmental potential. Whether similar mechanisms operate at other classes of chromatin bookmarks in pluripotent or multipotent stem cells remains an important question for future studies.

Our study also reveals a surprisingly complex relationship between KAT6 and KAT7. Both HAT families have attracted considerable attention in recent years due to their involvement in cancer^75–78^, their targetable nature and the promising results of combination therapy^41–43,79,80^. However, the molecular basis of the KAT6/7 synergies and compensations remains understudied. Our experiments uncovered unappreciated aspects of their genomic binding and activity, pointing towards both shared and unique behaviors. Despite their largely overlapping genomic occupancy, KAT6 and KAT7 proteins show different degrees of mitotic retention and different binding patterns. More specifically, their differential binding on promoter nucleosome-free regions suggests either distinct mechanisms of chromatin association/recruitment (through nucleosomes or other chromatin-bound factors) and/or potentially additional roles beyond catalyzing histone acetylation. The perturbation of mitotic chromatin composition (e.g. partial dissociation of mitotic SMARCB1) upon acute KAT6/7 depletion suggests a “scaffolding” function for these proteins or their acetylated targets. In terms of their catalytic activity, while KAT7 appears to be the dominant regulator of H3K14 acetylation, combined KAT6/7 perturbation produced substantially stronger phenotypes than disruption of either enzyme alone, supporting functional cooperation and partial redundancy. Moreover, the stronger transcriptional effects upon acute perturbation during M-to-G1 compared to prolonged KAT6/7 perturbation suggest a particularly important role of these proteins during mitotic exit and the existence of compensatory mechanisms that buffer core transcriptional programs in interphase. Together, our results underscore the high complexity of KAT regulation, interconnections and functions. Future individual and combinatorial perturbation studies employing catalytic inhibitors, mutants and degrons will be required to dissect cooperativities and redundancies, and distinguish between catalytic and non-catalytic effects.

The dual activity of KAT6/7 on both self-renewal and lineage-related genes may have implications across development and disease. In addition to the increasingly appreciated roles of KAT7 and KAT6 proteins in leukemia, breast cancer and other malignancies^41–43,78^, KAT6A and KAT6B mutations underlie a spectrum of neurodevelopmental disorders characterized by developmental delay, intellectual disability, and impaired neuronal maturation^44–50^. We speculate that different aspects of KAT-dependent mitotic chromatin regulation might underlie these distinct disease contexts. In development, KAT6/7-dependent chromatin states, i.e. the trivalent H3K14ac/H4me3/K27me3 state, may be particularly important for preserving the competence of stem and progenitor cells to activate lineage-specific programs at the correct developmental stage. In contrast, in cancer, KAT6/7 bookmarking may be co-opted to stabilize self-renewal networks and reinforce malignant cell identity across successive cell divisions, as has been reported for the KAT6/7-interacting chromatin modulator, MLL^18,42^. Understanding how KAT-dependent mitotic chromatin states balance cellular memory with developmental plasticity may therefore provide new insights into both normal development and disease pathogenesis.

Taken together, our findings support a model in which KAT6 and KAT7, actively maintain mitotic histone acetylation and chromatin composition both in mouse and human ESCs. Beyond preserving ongoing transcriptional states, KAT6/7 activity contributes to the priming of lineage-associated genes, thereby linking mitotic chromatin state to developmental competence. This expanded view of mitotic bookmarking provides a conceptual framework for understanding how stem cells balance the competing demands of cellular memory and developmental potential across successive cell divisions.

## Methods

### Mouse ESC culture

ESCs were cultured at 37°C on surfaces pre-coated with 0.2% w/v gelatin (Sigma-Aldrich, G1890). They were maintained in standard serum/2i/LIF conditions using DMEM (Gibco, 10829018) supplemented with 15% fetal bovine serum (R&D Systems, S10250), 100 U/mL penicillin-streptomycin (Gibco, 15140163), 1X non-essential amino acid (Gibco, 11140076), 1X GlutaMAX supplement (Gibco, 35050079), 55 µM β-mercaptoethanol (Gibco, 92185023), 128 µg/mL LIF (In-house), 1 μM MEK1/2 inhibitor (PD0325901) (Tocris, 4192) and 3 μM GSK3 inhibitor (CHIR99021) (Tocris, 4423) (from now referred to as 2i).

### Mouse ESC cell line construction

HaloTags® and dTAG cell lines were derived from Bruce-4 C57BL/6J mouse embryonic stem cells (ESCs; MilliporeSigma, CMTI-2) as previously established^84^. Homology arms covering 529 bp upstream and 652 bp downstream of the mKat6b and 1000 bp upstream and 1000 bp downstream of the mKat7 C-terminus were PCR amplified from genomic DNA and Topo cloned into a pCR2.1 plasmid (Thermo Fisher, 450641). A construct containing either the FKBP12F36V-2xHA-P2A-NLS-EGFP/mCherry or the 2xHA-HaloTag®-P2A-NLS-EGFP/mCherry was cloned into the pCR2.1 containing the homology arms by Gibson assembly (NEB, E5510S) ‘breaking’ the STOP codon. These targeting vectors were transfected together with a pX330-geneticin vector bearing the specific gRNAs targeting the C-terminus of both genes (mKat6B-GGCTGTCTACCTTCTCATGT; mKat7-TAGGTCACTTTAAGTGCCCT:) using TransIT-293 (Mirus Bio 2700) lipofection. To generate the dual mKat6B/mKat7 degron, cells already edited for mKat7 were transfected with the mKat6B targeting construct and gRNA. After transfection, cells were selected for the presence of pX330 with G418 (500 μg/ml) during 48 h. 2-3 days after selection cells expressing either the EGFP (mKat7) and/or the mCherry (mKat6B) transcriptional reporter, indicative of the correct genomic insertion, were single-cell sorted by FACS and expanded. Finally, individual clones were characterized by genotyping PCR, FACS to measure the levels of the transcriptional reporter and Western Blot to confirm homozygosity.

### Human ESC culture

Human MEL-1 ESCs were cultured on plates coated with vitronectin (Thermo Fisher, A14700) in Essential 8 Flex medium with supplements (Thermo Fisher, A2858501) and normocin (50 µg/ml) (InvivoGen, ant-nr-05). Cells were passaged with home-made EDTA dissociation solution (0.5 mM EDTA and 35 mM NaCl in 1x PBS) during maintenance or with Accutase (Thermo Fisher, 00-4555-56) for freezing or plating for experiments. Cell stocks were frozen in Stem Cell Banker (Amsbio, 11890). After thawing and single-cell seeding, ROCK inhibitor (Y-27632) (Tocris Bioscience, 27632) at 10 μM was added to the cell culture for 18-24 h.

### Human ESC cell line generation

*KAT7, KAT6A and KAT6B* HaloTags® and *KAT7*-dTAG hESCs were derived from human MEL-1 ESCs by CRISPR/Cas9 engineering as previously described^84^. Homology arms covering 697 bp upstream and 1000 bp downstream of *KAT7* C-terminus, 619 bp upstream and 1000 bp downstream of *KAT6A* C-terminus and 591 bp upstream and 563 bp downstream of *KAT6B* C-terminus were cloned/purchased into a 2xHA-HaloTag®-P2A-NLS-EGFP and additionally into a FKBP12F36V-2xHA-P2A-NLS-EGFP for *KAT7*. These targeting vectors were transfected together with a pX330-geneticin vector bearing the specific gRNAs targeting the C-terminus of each gene (hKAT7-gRNA, hKAT6B-gRNA) using lipofection in mTeSR media (Stem Cell Technologies, 05850) supplemented with CloneR (Stem Cell Technologies, 05888) solution (1:10). Cells were selected for the presence of pX330 with G418 for 48 h (500 μg/ml – Day 1, 250 μg/ml – Day 2). 2-3 days after selection cells expressing the EGFP transcriptional reporter were single-cell sorted by FACS and expanded for characterization. Homozygosity was assessed in the Individual clones by genotyping PCR and by WB when antibodies were available.

### Flow cytometry

Single cell-sorting of dTAG and HaloTag® clones was conducted with the help of the flow cytometry core facility at Weill Cornell. Live cells were run on the BD Influx, gated for high levels of mCherry and/or EGFP to enrich for homozygosity and seeded in 96-wellplates. After growing, single-cell derived clones were characterized by live-cell flow. Cells were resuspended in 3 µg/ml DAPI and filtered through a 35 µm cell strainer (VWR, 352235) into FACS tubes. Expression of endogenous loci bearing mCherry was determined using the 561 nm (610/20) channel while for EGFP the 488 nm (530/30) was used. Analysis was performed in a BD Fortessa flow cytometer and using the FlowJo v10.10 software.

To assess HA depletion after dTAG treatment, we assessed protein levels by intracellular flow cytometry. Briefly, 750,000 cells were collected in a single-cell suspension and processed using an intracellular staining kit (Thermo Fisher, Cat. 00-5523-00). Briefly, cells were fixed for 25 min followed by incubation with 100 µl of HA antibody diluted in permeabilization buffer (CST-3724S, 1:800) for 40 min. Cells were then incubated with 100 µl of secondary antibody (Thermo Fisher, A-21448, .1:500) for 30 min in the dark. Finally, cells were resuspended in 3 µg/ml DAPI and filtered through a 35 µm cell strainer (VWR, 352235) into FACS tubes for analysis. Analysis was performed in a BD Fortessa flow cytometer and using the FlowJo v10.10 software.

To quantify the surface marker SSEA-1, live cells were stained with anti-SSEA1-647 (Biolegend, 125608,1:600) for 20 minutes. In the final step, DAPI was added at 3 µg/ml DAPI. Analysis was performed using BD FACS Canto II analyzer and FlowJo v10.10 software.

### Western Blotting

Cells were lysed either in RIPA buffer (Thermo Fisher, 89900) with Halt protease inhibitors (Thermo Fisher, 1861280) for a total protein extract or with a histone extraction kit (Abcam, ab113476) for analysis of histone posttranslational modifications. Protein extracts were quantified with a detergent compatible protein assay (Bio-rad,5000111). Proteins were separated by SDS-PAGE using 4-12% polyacrylamide gels (Thermo Fisher, NP0322BOX) and in turn, transferred onto methanol activated PVDF membranes (Bio-rad, 1620177). The membranes were blocked in 5% non-fat milk (Santa Cruz, 2325) in TBS-T (0.5% Tween-20 (Millipore, 8170721000) in Tris Buffered Saline) overnight and subsequently incubated with the primary antibody of interest (reference table). As secondary antibodies, either IRDye® 800CW goat anti-rabbit IgG (Li-cor, P/N: 925-32214) or Alexa Fluor^TM^ 680 donkey anti-Mouse IgG (Thermo Fisher, A10038) were used. Blots were imaged and quantified using Image Studio Lite (Odyssey CLx imager, Li-COR Biosciences).

### Immunofluorescence

For immunofluorescence, cells were grown or attached to 18-mm round coverslips (Electron Microscopy Sciences, 72222-01). Cells were fixed by addition of 1% paraformaldehyde (Thermo Fisher, 033314.AP) for 10 minutes at room temperature. Cells were washed twice with phosphate-buffered saline (1x PBS) and then blocked and permeabilized in 0.5% bovine serum albumin (BSA) (Sigma, A3311-50G)) and 0.1% Triton X-100 (Sigma, X100-500ml) in PBS for 30 min at room temperature. Primary antibodies were incubated at specific concentration (reference table) in the same blocking buffer for 1 h at room temperature. Next, slides were washed thrice with blocking buffer and secondary antibodies conjugated with AlexaFluors were added in blocking buffer for 45 min at room temperature. Finally, slides were washed and Hoechst was added at 3 ug/ml or 3 min to stain DNA (Thermo Fisher, H3570). Coverslips were mounted on microscope slides with Prolong™ Gold Antidade Mountant (Thermo Fisher, P36930).

### Mouse ESC mitotic arrest and release

For mitotic collections cells were treated as previously described^85^. Briefly, mKat6B and Kat7-dTAG cells were plated as single-cells at a density of 45000 cells/cm^2^. After 6 hours, the cells were treated with 9 µM CDK1 inhibitor (Ro-3306; Selleckchem, S7747) for 16 hours for G1 synchronization. Next, the cells were washed twice with 1x PBS containing 100 ng/ml nocodazole (Sigma-Aldrich, M1404) and incubated in 2i media containing the same nocodazole concentration for 2 hours to block the cells in prometaphase. When relevant, DMSO or 500 nM dTAG-13 (Tocris, 6605) were added during those 2 hours for targeted protein degradation (i.e: Mitotic H3K14ac ChIP-seq upon dual-depletion). To enrich for the mitotic fraction, cells were subjected to “mitotic-shake-off” as mitotic cells are round and detach easier from the plates. Mitotic cells were washed thrice with PBS and their viability was quantified. Non-synchronized cells were plated and collected in parallel as a control. If required, cell pellets were snap-frozen and subsequently processed for chromatin, histone proteomics and WB analyses. If for ChIP-seq, cells were fixed as described in the ChIP-seq section. For release into G1, after mitotic shake-off cells were seeded on gelatin-coated plates while treated in DMSO/dTAG-13 for 1 h or 3h. Cells were snap-frozen at each timepoint for transcriptomics.

### Human ESC mitotic arrest and release

To collect mitotic cells, human MEL-I cells were plated with ROCKi. After 24 h, the cells were treated with 200 ng/ml nocodazole (Sigma-Aldrich, M1404) for 16 h for arrest in prometaphase. When relevant, DMSO or a combination of 250 nM dTAG-v1 (Tocris, 6914) and 100 nM of PF-9363 (MedChem, HY-132283) were added 2h prior to the mitotic collection. “Shake-off” was performed to enrich for the mitotic fraction and cells were washed thrice with 1x PBS before viability evaluation. Asynchronous cells were plated and collected in parallel. If needed, cell pellets were snap-frozen and subsequently processed for chromatin and histone proteomics. For release into G1, after mitotic shake-off cells were seeded on vitronectin-coated plates for 5 h while keeping the treatments. Cells were snap-frozen at that point for transcriptomics analysis.

### Validation of mitotic arrest and release

To validate the mitotic purity of cells, “shake-off” cells resuspended in PBS were seeded on poly-L-lysine (MilliporeSigma, P8920)-coated 18 mm round slides (Electron Microscopy Sciences, 72222-01). Cells were centrifuged at 1200 rpm for 2 min for attachment and subsequently fixed and processed as described in the immunofluorescence chapter for detection of the mitotic marker H3S10ph. For cell cycle analysis using fluorescence-activated cell sorting (FACS) analysis, 100,000-300,000 cells were resuspended in 1x PBS and fixed by adding 600 μl of ice-cold 100% ethanol while vortexing the tube at medium speed. Cells were kept on ice for at least 10 min and centrifuged at 500g for 5 min at 4 °C. The cell pellet was resuspended in PBS containing 0.3 μg/mL DAPI and filtered through at 35 μm cell strainer (VWR, 352235) into FACS tubes. The cell cycle was analyzed using a BD FACS Canto II analyzer and FlowJo v10.10 software.

### Neuroectoderm specification in hESCs

hESCs were differentiation into 10-day neuroectodermal cells as previously established^67^. ESCs were plated at a density of 3×10^5^ cells/cm^2^ in Essential 8 flex media with 10uM Y-27632 on Matrigel (354277, Corning) coated plates (d-1). On days 0-9 the culture media was replaced daily with Essential 6 medium (E6; Thermo Fisher Scientific, #A1516401) supplemented with 10 μM SB431542 (Tocris, #1614) and 100 nM LDN193189 (Stem Cell Technologies, #72142); with an additional 2 μM XAV939 (3748, Tocris) from days 0-2 to induce anterior neuroectodermal patterning.

### Histone proteomics

Histones were extracted from frozen pellets using a commercial histone extraction kit following the manufacturer’s protocol (Abcam, ab113476). In the subsequent steps, histones were processed as previously described by Sidoli et al^86^. Briefly, soluble histones were precipitated overnight with 33% cold trichloroacetic (TCA, Sigma-Aldrich, T0699-100ml) under rotation at 4°C. Samples were spun at 3400 rcf for 5 min, supernatant was removed and histone pellets were rinsed with ice-cold acetone + 0.1% HCl and with 100% ice-cold acetone. Histone pellets were allowed to dry at room temperature Next, the pellet was dissolved in 50 mM ammonium bicarbonate, pH 8.0, and histones were subjected to derivatization using 5 µL of propionic anhydride and 14 µL of ammonium hydroxide (all Sigma Aldrich) to balance the pH at 8.0. The mixture was incubated for 15 min and the procedure was repeated. Histones were then digested with 1 µg of sequencing grade trypsin (Promega) diluted in 50mM ammonium bicarbonate (1:20, enzyme:sample) overnight at room temperature. Derivatization reaction was repeated to derivatize peptide N-termini. The samples were dried in a vacuum centrifuge. Prior to mass spectrometry analysis, samples were desalted using a 96-well plate filter (Orochem) packed with 1 mg of Oasis HLB C-18 resin (Waters). Briefly, the samples were resuspended in 100 µl of 0.1% TFA and loaded onto the HLB resin, which was previously equilibrated using 100 µl of the same buffer. After washing with 100 µl of 0.1% TFA, the samples were eluted with a buffer containing 70 µl of 60% acetonitrile and 0.1% TFA and then dried in a vacuum centrifuge.

Samples were resuspended in 10 µl of 0.1% TFA and loaded onto a Dionex RSLC Ultimate 300 (Thermo Scientific), coupled online with an Orbitrap Fusion Lumos (Thermo Scientific). Chromatographic separation was performed with a two-column system, consisting of a C-18 trap cartridge (300 µm ID, 5 mm length) and a picofrit analytical column (75 µm ID, 25 cm length) packed in-house with reversed-phase Repro-Sil Pur C18-AQ 3 µm resin. Peptides were separated using a 30 min gradient from 1-30% buffer B (buffer A: 0.1% formic acid, buffer B: 80% acetonitrile + 0.1% formic acid) at a flow rate of 300 nl/min. The mass spectrometer was set to acquire spectra in a data-independent acquisition (DIA) mode. Briefly, the full MS scan was set to 300-1100 m/z in the orbitrap with a resolution of 120,000 (at 200 m/z) and an AGC target of 5×10e5. MS/MS was performed in the orbitrap with sequential isolation windows of 50 m/z with an AGC target of 2×10e5 and an HCD collision energy of 30.

Histone peptides raw files were imported into EpiProfile 2.0 software^87^. From the extracted ion chromatogram, the area under the curve was obtained and used to estimate the abundance of each peptide. In order to achieve the relative abundance of post-translational modifications (PTMs), the sum of all different modified forms of a histone peptide was considered as 100% and the area of the particular peptide was divided by the total area for that histone peptide in all of its modified forms. The relative ratio of two isobaric forms was estimated by averaging the ratio for each fragment ion with different mass between the two species. The resulting peptide lists generated by EpiProfile were exported to Microsoft Excel and further processed for a detailed analysis.

### Chromatin proteomics

For the differential asynchronous/mitotic proteomics, a commercial fractionation kit (Life Technologies, 78840) was used to extract the chromatin fraction of mitotic and asynchronous mESC and hESCs. Chromatin extracts were processed for proteomics as follows. Samples were reduced using 10 mM TCEP and 1% SDS for 1 h at 55 °C. Samples were then alkylated with 375 mM iodoacetamide for 30 min at room temperature in the dark. Trypsin Gold, Mass Spectrometry Grade (catalog no. V5280; Promega), was used for protein digestion at an enzyme substrate ratio of 1:100. The samples were incubated overnight at 37 °C. Peptide concentrations were quantified using the Pierce Quantitative Colorimetric Peptide Assay (catalog no. 23275; Thermo Scientific). For each sample, 500 ng of peptides was quenched by adding TFA and FA to acidify the solution. Peptides were desalted using a homemade spin column and resuspended in 10 μL of Solution A (0.1% formic acid in water). Aroud 500ng was injected for mass spectrometry analysis. The digested peptides were analyzed by nanoLC-ESI-MS/MS using a timsTOF HT (Bruker) coupled to a nanoElute2 ultra-high-performance liquid chromatography (UHPLC) system. Peptide separation was performed on the nanoElute2 UHPLC system using a homemade C18 analytical column (100 μm inner diameter, 10 cm length, 1.9 μm C18 particle size) at a flow rate of 500 nL/min. Mobile phases consisted of Solution A (0.1% formic acid in water) and Solution B (0.1% formic acid in acetonitrile). The gradient was programmed as follows: 2% to 35% Solution B over 25 min, 35% to 95% Solution B over 0.5 min, followed by a 4.5 min hold at 95% Solution B. Prior to separation on the homemade analytical column, samples were loaded onto a Thermo PepMap™ Neo Trap Cartridge with PepMap™ Neo UHPLC Columns (Thermo Scientific, Cat# 174500). DIA data acquisition on the timsTOF HT was performed using Compass Hystar version 6.3 (Bruker) and timsControl version 6.0.2 (Bruker). The DIA-PASEF method covered an m/z range of 100–1700, with 60 windows spanning 300–1300 Da and an ion mobility range (1/K₀) of 0.60–1.45 V·s/cm². The ramp time was 75.0 ms, and the accumulation time was 50.0 ms. The capillary voltage of the CaptiveSpray source was set to 1500 V, with a dry gas temperature of 180 °C and a dry gas flow rate of 3.0 L/min. The collision energy was set to 10.0 eV. Before each set of runs, m/z and ion mobility calibration was performed using three reference ions from the ESI-Filter Cal (m/z 622, 922, and 1222). DIA data were analyzed using DIA-NN (version 1.8.1) with the Bruker mouse proteome spectral library (Bruker_Mouse_Trypsin_TIMScore_v2.tsv). The analysis parameters included up to three missed cleavages, a maximum of two variable modifications, a precursor charge range of 1–4, and a precursor m/z range of 100–1800. Data were processed in double-pass mode for the neural network classifier, with “Any LC (high accuracy)” selected for quantification. Match Between Runs (MBR) was enabled. All other settings were kept at their default values, with RT-dependent cross-run normalization applied and results filtered at a 1% FDR. The analysis was performed using 32 threads, as automatically recommended by the software.

### Live-cell imaging

For live-cell imaging mouse ES cells were grown on Biolaminin (Biolamina, LN511-0202)-coated chambers (Ibidi, 80826) 24 h prior to imaging. Human cells were grown on vitronectin-coated 35 mm dishes (Maltek, P35-1.5-14-C.s) 24h prior to imaging. Cells were labeled with 200 nM JF549 (Promega, GA1110) or 200 nM J503 (Promega, HT1010) and 3 μM SiR-DNA (SpiroChrome, SC0007) for 1 h at 37 °C and washed prior to imaging. Cells were imaged on a Nikon Ti2-E motorized inverted microscope equipped with a Yokogawa CSU-W1 SoRa spinning-disk confocal unit and a Hamamatsu Orca Fusion BT sCMOS camera, using a 60× Apo TIRF oil-immersion objective (NA 1.49). Cells were maintained at 37 °C and 5% CO2 in a Tokai Hit stage-top incubator. For each field, a z-stack of 7 planes was acquired at 0.2 µm axial steps. Acquisition was controlled with NIS Elements. Chromatin enrichment quantification was performed as previously described^88^. Briefly, ImageJ was used to measure the mean HaloTag® fluorescence intensity in the area of the mitotic chromatin (selected from the SiR-DNA stain) and the mean cytoplasmic HaloTag® intensity. The chromatin enrichment score was calculated by taking the log2 ratio of the chromatin mean/cytoplasmic mean.

### ChIP-seq

ChIP-seq was performed as previously described (https://pmc.ncbi.nlm.nih.gov/articles/PMC7339746/). For histone PTM mapping, mESC and hESCs cells were crosslinked in 1 % w/v formaldehyde at room temperature for 10 min and quenched with 125 mM glycine for 5 min at room temperature. Crosslinked cells were centrifuged at 1,250rcf for 5 min at 4 °C and were washed twice with ice-cold PBS before snap-freezing. To map binding of mKAT7, mKAT6B, hKAT7 and hKAT6A, cells were subjected to a double-crosslinking protocol as previously described^89^. Cells were crosslinked with DSG (2 mM) (Sigma, 80424-5 mg) for 50 min at room temperature followed by 10 min with 1 % w/v formaldehyde (Thermo Fisher, 28908). 125 mM glycine was added for 5 min for quenching and cells were washed with ice-cold 1x PBS prior to snap-freezing. To quantify relative abundances, we added 5% spike-in of 1% or double-fixed MEL-I cells to mESCs or the opposite combination in the case of hESCs. Cells were used for mKAT7 and mKAT6B ChIP (10-15 x 10^6^), H3K14ac and H3K23ac ChIP in mESCs (2-5 x 10^6^), hKAT7 and hKAT6A ChIP (30 x 10^6^) and human H3K14ac ChIP (5 x 10^6^). Cells were lysed in lysis buffer (10 mM Tris pH 8, 1 mM EDTA (Thermo Fisher, AM9260G) and 0.5% SDS (Sigma, 05030-500ml-F) supplemented with protease inhibitors and incubated on ice for 20 min. The cells were sonicated in a Bioruptor device (30 s on/off; high setting) and spun down at the maximum speed for 15 min at 4 °C. The supernatants were pre-cleared under rotation for 1h at 4 °C with protein G Dynabeads (Invitrogen, 10004D) pre-blocked with BSA-protein (Sigma, 9048-46-8). Next, the supernatants were diluted five times with dilution buffer (0.01% SDS, 1.1% Triton X-100, 1.2 mM EDTA, 16.7 mM Tris pH 8 and 167 mM NaCl) and incubated overnight with the respective antibody with rotation at 4 °C. Protein G Dynabeads pre-blocked with BSA protein were added the following day and incubated for 3-4h at 4 °C. The beads were immobilized on a magnet and washed twice with low-salt buffer (0.1% SDS, 1% Triton X-100, 2 mM EDTA, 150 mM NaCl and 20 mM Tris pH 8), twice in high-salt buffer (0.1% SDS, 1% Triton X-100, 2 mM EDTA, 500 mM NaCl and 20 mM Tris pH 8), twice in LiCl buffer (0.25 M LiCl, 1% NP-40, 1% deoxycholic acid (sodium salt), 1 mM EDTA and 10 mM Tris pH 8) and once in TE buffer. DNA elution from the beads was done by incubation with 125 µl elution buffer (1% SDS and 10 mM NaHCO_3_) for 30 min at 65 °C 2000 rpm in a thermomixer. The supernatants and the inputs were reverse crosslinked by incubation overnight at 65 °C with 100 µg/ml amount of proteinase K. After treatment with RNase A for 30 min at 37 °C, the DNA was purified using a kit for DNA cleaning and concentration (Zymo, zd4014). The immunoprecipitated DNA was used for library preparation using the KAPA Hyper prep kit (KAPA Biosystems, 7962347001). The libraries were sequenced on an Illumina NovaSeq X plus 1.5B/10B Flow Cell (PE100).

### RNA-seq

Total RNA was extracted from 500,000 cells per sample using RNeasy Mini Kit (Qiagen, 74104), following the manufacturer’s instructions. Libraries were prepared by Novogene using their mRNA library preparation service with poly A enrichment. Libraries were sequenced on Illumina NovoSeq X Plus in PE150 mode.

### RT-qPCR

For RT-qPCR, total RNA was extracted from 50,000 cells in G1 using RNeasy Mini Kit (Qiagen, 74104). 500 ng of total RNA were retrotranscribed into cDNA using the iScript kit (Bio-Rad, 1708841). qPCR was performed in triplicates using PowerUp SYBR green PCR master mix (Thermo Fisher A25778) on an Applied Biosystems QuantStudio3. Primers spanning exon-intron overlaps are shown in Suppl. Table 7.

### ATAC-seq

ATAC-seq was performed following previously published protocols^90^. After mitotic shake-off, 50,000 cells per replicate were washed in 1x PBS and resuspended in 50 µl ATAC-resuspension buffer (10 mM Tris-HCl pH 7.4, 10 mM NaCl, 3 mM MgCl_2_) supplemented with 0.1% (v/v) IGEPAL CA-630, 0.1% Tween-20, 0.01% Digitonin for nuclei isolation. Samples were incubated on ice for 3 min, spun down and washed with 1 ml of cold ATAC-resuspension buffer containing 0.1% Tween-20. Nuclei were pelleted at 500 rcf for 10 min at 4 °C. After careful aspiration of the supernatant, the cell pellet was resuspended in 50 µl of transposition mix (25 µl TD buffer, 0.8 ul homemade Tn5, 18.2 µl 1x PBS, 0.5 µl 1% digitonin, 0.5 µl 10% Tween-20, 5 µl H_2_O) by pipetting up and down 6 times. The reaction was incubated at 37 °C for 30 min in a thermomixer at 1000 rpm. DNA was isolated using a commercial kit (Zymo, zd4014). Libraries were generated using the NEBNext HiFi 2x PCR master mix (NEB, #M0541) with each sample having a unique barcode and a universal primer by 10 cycles amplification. Finally, samples underwent a double-size selection (0.5x-1.3x) with SPRIselect beads (Beckman Coulter, B23317) and sent for Tapestation quality check. Librariers were sequenced on an Illumina NovaSeq X plus 1.5B/10B Flow Cell (PE50).

### Quantification and statistical analysis

Sequencing data (RNA-seq and ChIP-seq) and statistical analyses were processed using the previously described analysis pipeline^85^. Any modifications are described in the corresponding sections below. Snakemake (v7.32.4) was used to run pipelines for all genomic analyses^91^ All statistical analyses and figure generation in this manuscript were done using R (v4.4.0). Two-tailed Wilcoxon rank-sum test was applied to compare distributions between two groups after confirming that the data did not follow a normal distribution. Fisher’s exact test was used to evaluate enrichment of a variable of interest between the two groups. Additionally, K-means clustering was used to identify groups of genes with similar expression patterns across the dataset.

### RNA-seq analysis

Differential analysis was performed with R package DESeq2 (v1.44.0)^92^ using an adjusted p-value of < 0.05 as a cutoff. Raw read counts, DESeq2 normalized read counts, and DESeq2 analysis are provided in Suppl. Table 5.

### ATAC-seq analysis

Adapters were trimmed off from paired-end reads using bbduk.sh (v2.4.5)^93^ with the following option: ktrim=r ref=adapters rcomp=t tpe=t tbo=t hdist=1 mink=11. Filtered reads were aligned to the mouse genome (mm10) using bowtie2 (v2.4.5)^94^ with the options --end-to-end --very-sensitive --no-mixed --no-discordant --no-unal -X 2000. Mapped primary alignments with MAPQ ≥ 20 were retained using samtools with the options -q 20 -F 4 -F 256. PCR duplicates were removed using the MarkDuplicates module from picard (v2.27.4) (https://broadinstitute.github.io/picard/) with ^86^the following options: VALIDATION_STRINGENCY=LENIENT REMOVE_DUPLICATES=true. Mitochondrial reads and reads overlapping mm10 blacklist regions were removed using samtools and BEDTools (v2.30.0)^93^. BEDTools genomecov was used to convert final BAM files to bedGraph format with read-depth normalization to one million mapped reads, and bedGraphToBigWig from kentUtils was used to convert bedGraph files to bigWig format. Peaks were called with MACS2 callpeak.

### ChIP enrichment analysis and gene ontology analysis

ChIP enrichment analysis of ChIP peaks was performed using LOLA (v1.34.0)^83^. Only cell-type-specific ChIP-seq data from the LOLA reference database were used for enrichment analysis. In cases where multiple significant hits were returned for the same target, the result with the highest rank was selected as representative. The full list of ChIP enrichment analysis results is provided in Suppl. Table 4. Gene ontology analysis was performed using the enrichGO function in clusterProfiler (v.4.12.6)^95^. Benjamini-Hochberg correction with a p-value cutoff of 0.05 and a q-value cutoff of 0.2 was used. The full list of gene ontology analysis results is provided in Suppl. Table 4.

### Chromatin proteomics analysis

Protein-level quantitative outputs were generated using DIA-NN (v1.8.1). Downstream differential expression analysis was performed using MAGMa, a customized linear modeling framework based on the limma^96^ and MSstatsTMT^97^ R packages, following approaches previously described for differential analysis of mass spectrometry-based proteomics data. Briefly, protein abundance differences between experimental conditions were quantified by estimating fold changes and their associated statistical significance using moderated linear models. The core analytical framework and statistical modeling strategy have been described previously^98^, with internal extensions implemented here to support DIA-based quantification. For DIA-specific handling of missing values, imputation was performed under a missing-not-at-random assumption. Specifically, proteins missing across all replicates within a given condition were imputed using the minimum observed intensity for that protein within the corresponding condition. This approach assumes left censoring due to the limit of detection and leverages the high quantitative consistency of DIA data, in which systematic absence across replicates is interpreted as biologically meaningful signal loss rather than stochastic dropout. Proteins were considered significantly differentially abundant if they exhibited a fold change greater than 2 and a Benjamini–Hochberg-adjusted false discovery rate (FDR) of less than 0.05, following differential analysis.

## Data availability

All genomic datasets generated in this study, including RNA-seq, ChIP-seq, ChIP-exo, and ATAC-seq, have been deposited in the Gene Expression Omnibus (GEO) under accession number GSEXXXXX. Quality control metrics for genomics data, differential analysis results from RNA-seq, ChIP enrichment analysis, gene ontology analysis, K-means clustering are provided in Suppl Table 6. The mass spectrometry proteomics data have been deposited to the ProteomeXchange Consortium via the PRIDE^99^ partner repository with the dataset identifiers PXDXXXXX and PXDXXXXX.

## Acknowledgments

We sincerely thank all members of the Apostolou and Stadtfeld groups for their critical feedback throughout this study. We especially appreciate technical support from Kaushiki Chatterjee and Emily Goetlzer. U.L. was supported by the Korea Foundation for Advanced Studies (KFAS) Overseas PhD Scholarship. B.J.K. was supported by a Medical Scientist Training Program grant to the Weill Cornell/Rockefeller/Sloan Kettering Tri-Institutional MD-PhD Program (T32GM152349) from the National Institutes of Health. This study was supported by the Tri-Institutional Stem Cell Initiative (TriSCI), the KAT6 foundation and by the National Institute of General Medical Sciences (NIGMS) (RM1GM139738, R01NS136475 and R01GM138635). The Sidoli lab gratefully acknowledges funding from the Aging Biology Foundation, the Hevolution Foundation (AFAR), the ERCM-CFAR Center for AIDS Research, the NIAID for the PROVIDENT grant (1U19AI181977), the Einstein-Mount Sinai Diabetes center, and the NIH Office of the Director (S10OD030286).

## Author Contributions

E.A. conceived, designed, and supervised the study with input from A.L-D. A.L-D. performed all experiments in the study. U.L. led the computational analysis, with input from E.A. and A.L-D, and assisted with mitotic release and FACS experiments. X.L performed computational analysis related to mitotic bookmarking. A.L assisted with genotyping, cloning and Western Blot analyses. K.K helped with intracellular FACS and neuronal differentiation in hESCs. U.C.R. assisted with live-cell imaging microscopy with support by P.G. J.J.K and H.Y performed chromatin proteomics. S.S and S.S performed histone proteomics. B.J.K generated mouse KAT6B-Halo lines for live-cell imaging. S.K. assisted with immunofluorescences. M.S. helped with cloning strategies and provided scientific feedback throughout the project. A.L-D prepared all figures and, together with E.A, wrote the manuscript with input from all authors.

## Declaration of AI use

AI tools were used only for proofreading.

---

**Supp. Figure 1. Related to Figure 1.**
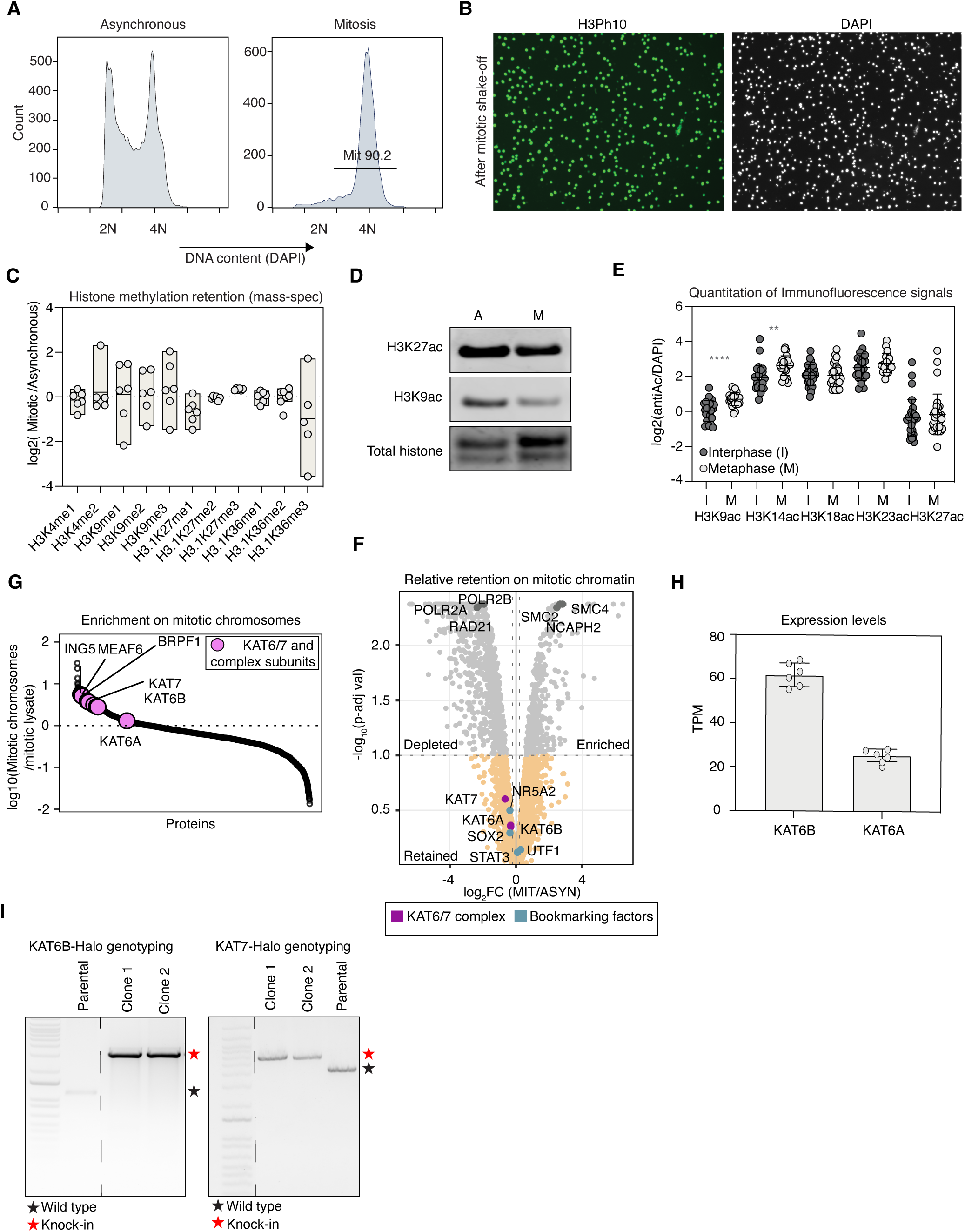
**A.** Representative FACs analysis of asynchronous (left) and mitotic mouse ESCs (after dual CDK1/nocodazole treatment and mitotic shake off), as assessed by measuring DNA content through DAPI staining. **B.** Representative immunofluorescence images showing the high purity of mitotic mouse ESCs after synchronization and shake-off. The mitotic marker, H3Ph10 (Histone-3-phosphoserine 10) is shown in green and DAPI in grey. **C.** Box plot showing the relative abundance of different histone methylation marks on mitotic and asynchronous histone extracts (expressed as log_2_ ratio) as detected by mass-spectrometry. Data from n = 6 biological replicates is shown as a mean. No statistical differences were found upon one-sample two-tailed t-test against 0. p > 0.05. The full list of histone proteomics data is provided in Suppl. Table 1. **D.** Western blot analysis showing asynchronous and mitotic levels of H3K27ac and H3K9ac, which were not detected consistently in our unbiased histone proteomics screening. Total histone was used as a loading control. Representative of n = 2 biological replicates. **E.** Dot plot depicting the quantification of the fluorescence intensity (Histone acetylation levels) across interphase and metaphase cells for both H3ac and H3K14ac. Fluorescence intensity was calculated as the mean signal of the Ac-histones on metaphase plates and interphase nuclei normalized by background subtraction. Statistical significance was assessed by two-tailed t-test comparing the mean interphase and metaphase levels. ** > 0.01, **** > 0.0001. (H3K9ac n= 22 cells, H3K14ac n= 23 cells, H3K18ac n= 28 cells, H3K23ac n= 22 cells, H3K27ac n= 29 cells). **F.** Volcano plot related to Figure 1F showing the enrichment or depletion of proteins on mitotic chromatin as determined by quantitative mass spectrometry. The log_2_fold change (mitotic/asynchronous) is represented on the x-axis and the -log_10_(p-adj val) on the y-axis. Mitotically enriched or depleted proteins (p-adj val <0.1) are shown in grey with control complexes highlighted in darker grey dots (p-adj val > 0.1). Retained proteins are depicted in orange (p-adj val > 0.1), while members of the KAT6/7 complex are in magenta, and previously characterized bookmarking factors are in blue. The full list of proteins related to this volcano plot is provided in Suppl. Table 2. **G.** Rank plot of proteins based on their relative abundance on purified mitotic chromosomes compared to mitotic lysates without chromosomes (expressed as log_10_ ratio) from published mass-spectrometry data^55^. Our candidate HATs and relevant subunits from their respective complexes are highlighted among the top enriched proteins on purified mitotic chromosomes. **H.** Bar plot showing the RNA expression levels (TPM) of *Kat6b* and *Kat6a* genes in our culture conditions in mESCs. Data from n = 6 biological replicates is shown. **i.** Representative genotyping PCRs to verify successful, homozygous knock-in of the HaloTag cassette at the endogenous *Kat6b* and *Kat7* genomic loci by CRISPR engineering compared to the wild-type bands in the parental lines. Primers are provided in Suppl. Table 7.

**Supp. Figure 2. Related to Figure 2.**
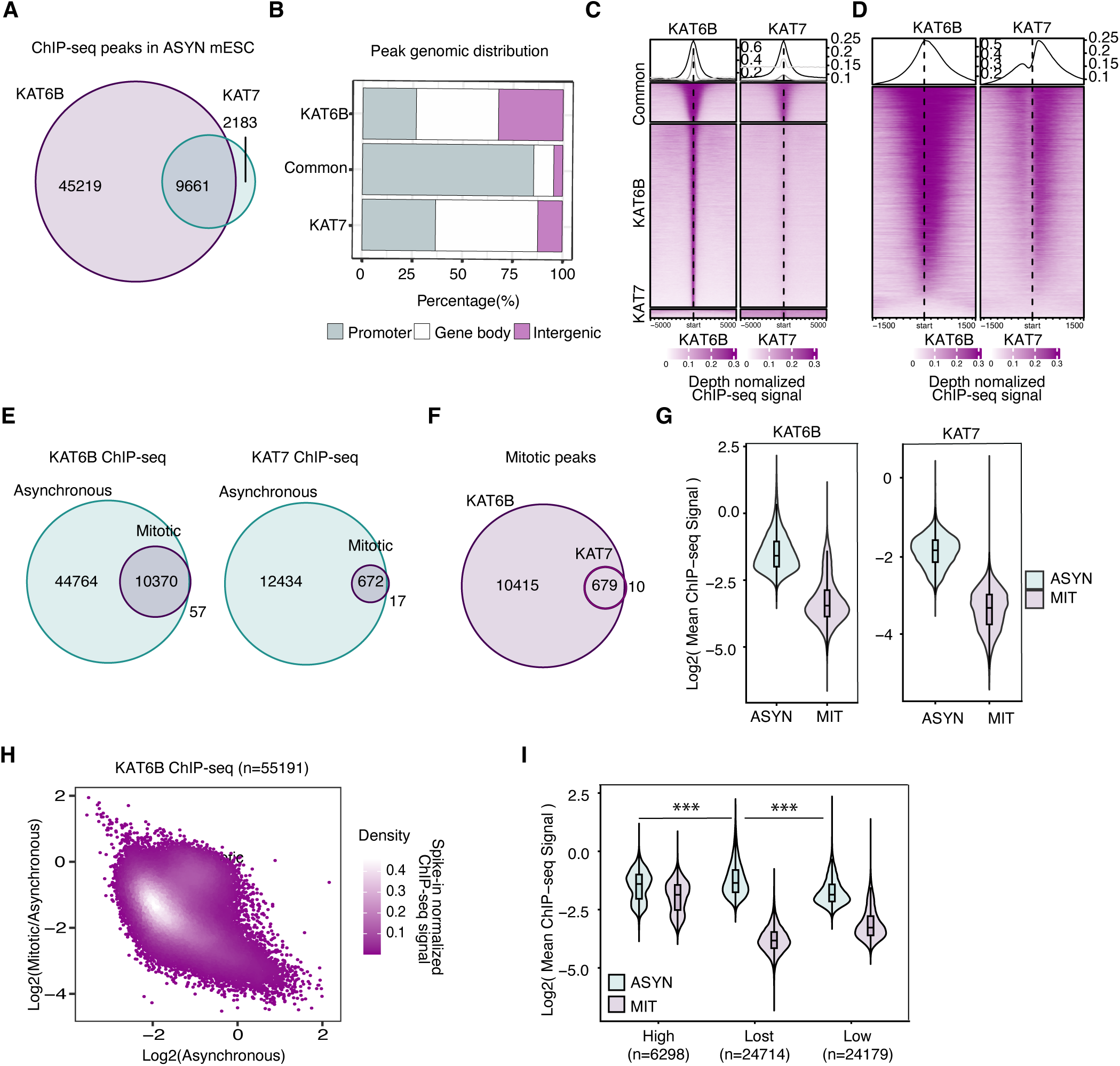
**A.** Venn diagram showing the overlap between the KAT6B and KAT7 ChIP-seq peaks in asynchronous mESCs. **B.** Stacked bar plot representing the genomic distribution (percentages) of the KAT6B-specific, KAT7-specific and common peaks from **Supp.** Fig.2A across promoters, gene bodies or intergenic regions. **C.** Tornado plots comparing the KAT6B and KAT7 ChIP-seq signal (normalized over input and sequencing depth) across KAT6B- and KAT7-specific as well as common peaks as assigned in **Supp.** Fig.2A. The average signal of n= 2 independent biological replicates is projected +/- 5 kb around peak centers. **D.** Tornado plots showing the KAT6B and KAT7 ChIP-seq signal (normalized over input and sequencing depth) around KAT6B-target genes. The average signal of n= 2 independent biological replicates is projected +/- 1.5kb around the Transcriptional Start Site (TSS). **E.** Venn diagram showing the degree of overlap between asynchronous and mitotic ChIP-seq peaks for KAT6B and KAT7. **F.** Violin plot showing the log_2_ mean ChIP-seq signal (normalized over input and sequencing depth) in asynchronous and mitotic cells for KAT6B and KAT7. **G.** Scatterplot comparing the relative change in KAT6B binding during mitosis over the asynchronous signal, as shown in Fig.2A, but this time the ChIP-seq signals are normalized over input and over spike-in control (instead of sequencing depth). **H.** Violin plot showing the log_2_ mean ChIP-seq signal in asynchronous and mitotic cells for KAT6B in the high, lost and low categories defined in Fig.2a. Two-sided Wilcoxon rank-sum test on asynchronous signals of pairs of the High, Low, Lost groups: Lost-Low (Effect size r=0.3848, P=0); Lost-High (Effect size r=0.1119, P=1.96×10^-86); Low-High (Effect size r=0.1709, P=1.24×10^-195).

**Supp. Figure 3. Related to Figure 3.**
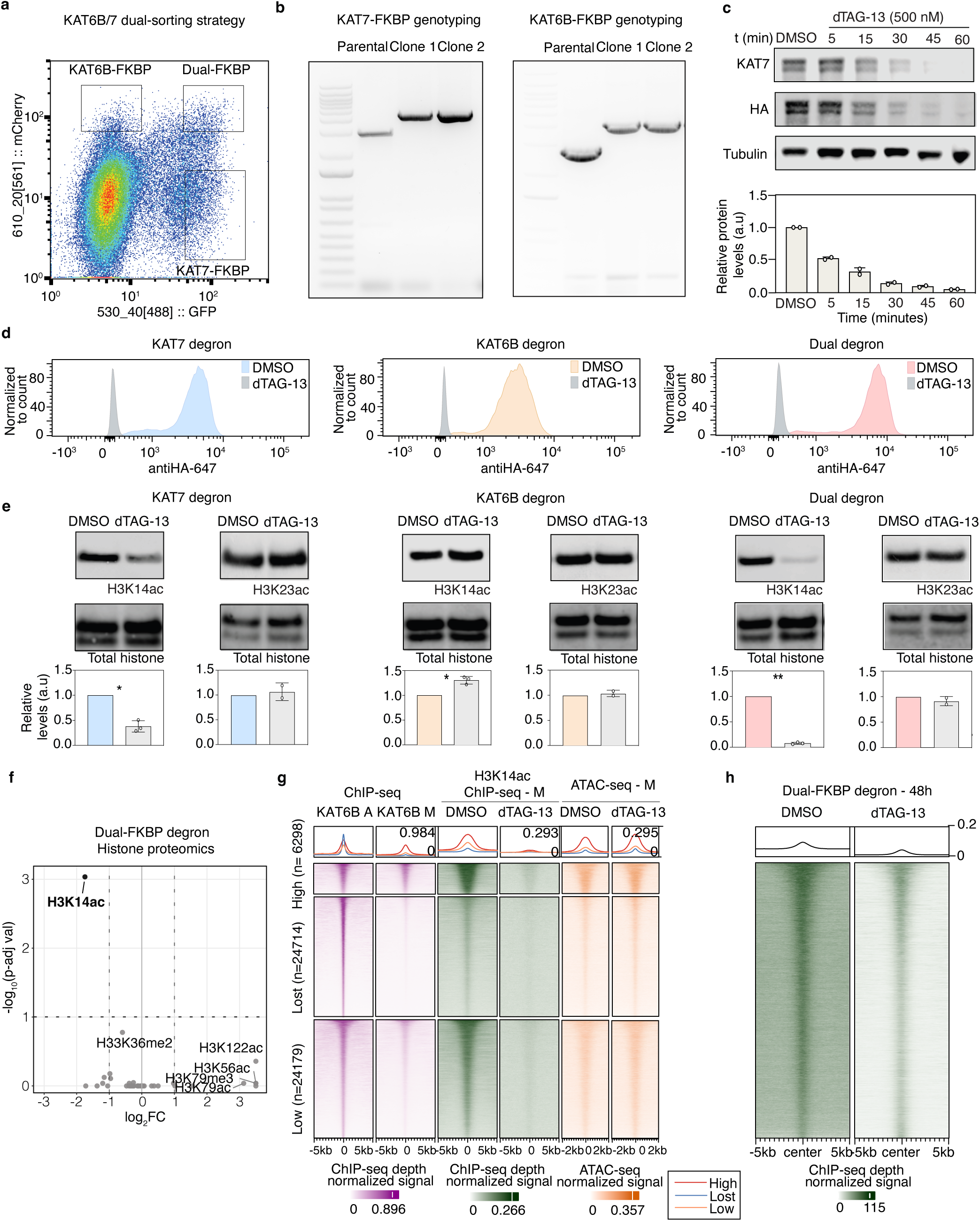
**A.** Representative FACs scatterplot depicting the gating strategy for sorting cells with successful integration of KAT6B-and KAT7-FKBP cassettes, individually or combined. GFP fluorescence (linked to the KAT7 cassette) is on the x-axis while mCherry (linked to the KAT6B cassette) is on the y-axis. Rectangles show the gating choices for generation of the different knock-in cell lines. **B.** Representative genotyping PCRs to verify successful, homozygous knock-in of the FKBP degron cassettes at the endogenous *Kat6b* and *Kat7* genomic loci by CRISPR engineering compared to the wild-type bands of the parental cells. For Dual-FKBP degrons homozygosity was validated for both genomic loci. Primers are provided in Suppl. Table 7. **C.** *Top:* Western blot analysis showing the depletion kinetics of the engineered KAT7-FKBP protein upon treatment with dTAG-13. Antibodies against KAT7 and against the HA-tag of the knock-in cassette were used. Tubulin served as a loading control. *Bottom:* Bar plot quantification of the Western blot results shown on the top. **D.** Histograms showing the successful KAT7 and KAT6B protein depletion by intracellular flow. Individual and dual-KAT6B/7 degrons were treated for 48 h with DMSO/dTAG-13 and the HA levels in both conditions were assessed by intracellular FACS. Cell-lines are color-coded. KAT7 DMSO is blue, KAT6B DMSO is orange and the dual-degron DMSO is pink. HA levels upon dTAG-13 treatment are grey for all cell-lines. The histograms are representative of n = 2 biological replicates per cell-line. **E.** Analysis of H3K14ac and H3K23ac levels after KAT7 and/or KAT6B protein depletion. *Top:* Representative Western blot analysis of histone extracts using anti-H3K14ac and anti-H3K23ac antibodies comparing their levels after 48 h depletion of KAT7 (blue), KAT6B (orange) or both (pink) upon dTAG-13 treatment. The total histone staining was used as a loading control. *Bottom:* bar plot quantification of H3K14ac and H3K23ac levels normalized to the histone loading and relative to the DMSO control. n= 2 biological replicates for H3K14ac. H3K14ac levels were significantly lower for KAT7 (*p < 0.05) and the dual-degron (**p < 0.05) after one-sample two-tailed t-test. H3K14ac levels were significantly higher for KAT6B (*p < 0.05). H3K23ac levels did not show significant differences. **F.** Volcano plot showing the differential analysis of histone posttranslational modifications, as detected by histone proteomics, after 48 h of KAT7 and KAT6B co-depletion. x-axis shows the log_2_ fold change (DMSO/dTAG-13) while the y-axis represents the -log_10_(p-adj val). H3K14ac is the only significantly reduced histone PTM upon co-depletion (log_2_FC > -1 and p-adj val< 0.1). The full analysis related to this volcano plot is provided in the Suppl. Table 1. **G.** Tornado plot showing the changes in mitotic H3K14ac ChIP-seq and ATAC-seq signals upon KAT6/7 co-depletion around the different KAT6B peak categories as defined in Fig.2A based on their degree of retention during mitosis (high, low, lost). The mitotic H3K14ac ChIP-seq (green, +/- 5kb peaks) and ATAC-seq (orange, +/- 2kb peaks) signals in DMSO and dTAG conditions are projected centered around the KAT6B peaks. Average depth normalized signal is shown for n = 2 biological replicates for KAT6B ChIP-seq and n = 2 technical replicates for H3K14ac ChIP-seq and ATAC-seq. **h.** Tornado plot showing H3K14ac ChIP-seq signal (normalized over sequencing depth) in asynchronous mESCs after 48 h of KAT6B/7 co-depletion in asynchronous mESCs (dTAG-13) in comparison with DMSO control. Average signal of n = 2 biological replicates is displayed.

**Supp. Figure 4. Related to Figure 4.**
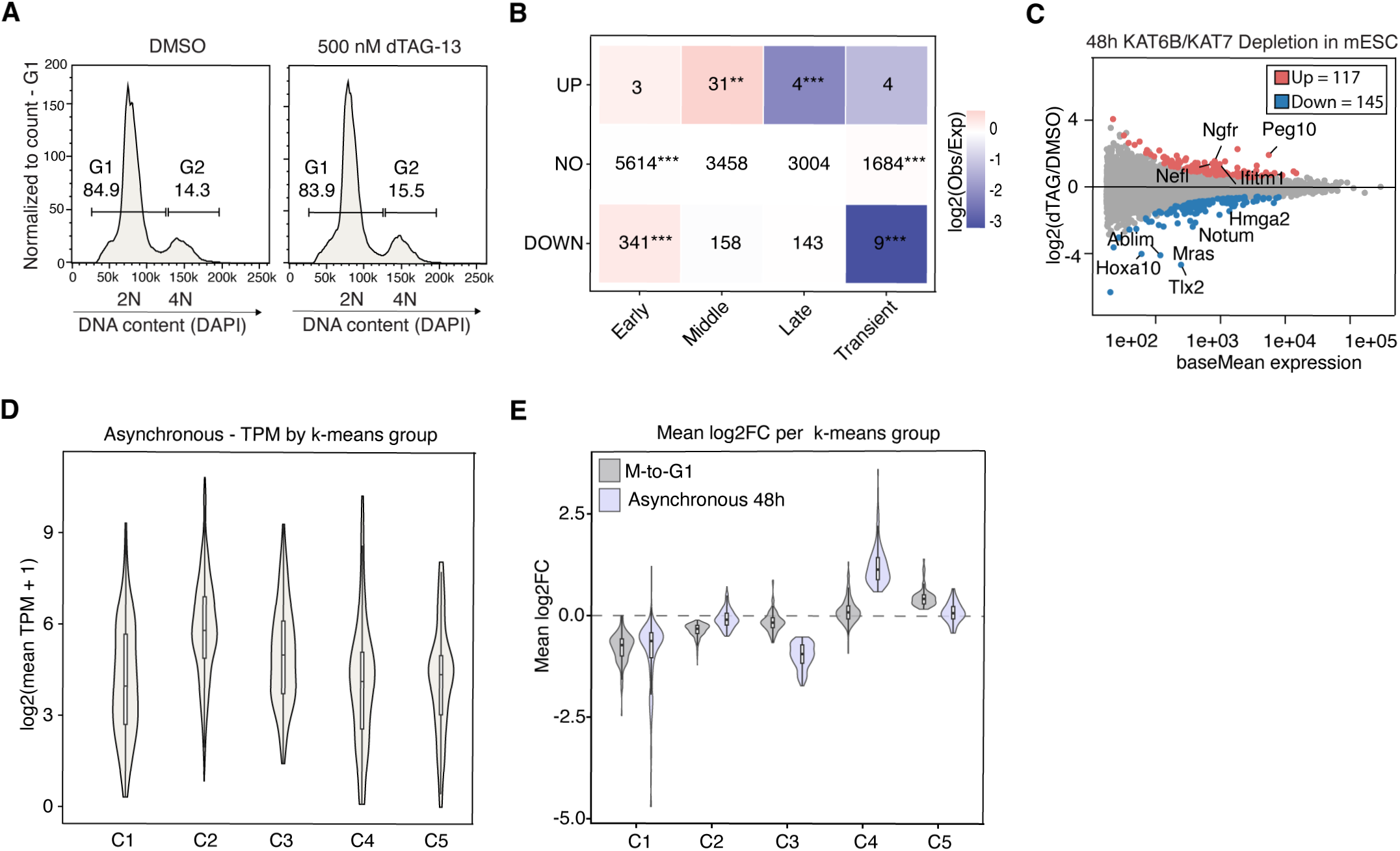
**A.** Cell cycle profile analyzed by FACS measuring DAPI (DNA) content after mitotic arrest and release for 3 h into G1 in DMSO and dTAG-13 treated cells. A prominent 2N peak (> 84%) of similar magnitude was consistently detected in control and dTAG-treated cells, indicating no cell-cycle defects in the release. **B.** Heatmap showing the relative enrichment of differentially expressed genes in G1 (Up, non-DEGs, down) upon KAT6/7 co-depletion for previously characterized gene groups with distinct postmitotic activation kinetics: early, mid, late and transient, as defined in^12^. The cell values indicate the number of genes in each condition and the color represents the log_2_(odds ratio) of enrichment or depletion, with red indicating over-representation and blue indicating under-representation relative to expected values. Statistical significance was assessed by Fisheŕs exact test (*< 0.05, **< 0.01, ***< 0.001). **C.** M/A plot of differentially expressed genes in mESCs upon KAT6B/7 co-depletion for 48hr (dTAG) versus DMSO. The log_2_ fold change (dTAG13/DMSO) is plotted on the y-axis against the average expression levels for each gene (baseMean) on the x-axis for DMSO (n = 2 technical replicates). p-adj val > 0.05 was uses as a cut-off to call differential expression. Each point represents a single gene. Up-regulated genes are colored in red (N = 117) and downregulated genes are colored in blue (N=145). Non-DEGs are grey. Names of representative genes are depicted. The full list of DEGs is shown in Suppl.Table 5. **D.** Violin plot showing the mean and distribution of the absolute expression levels per gene cluster as defined by the k-means clustering shown in Fig.4F.

**Supp. Figure 5 Related to Figure 5.**
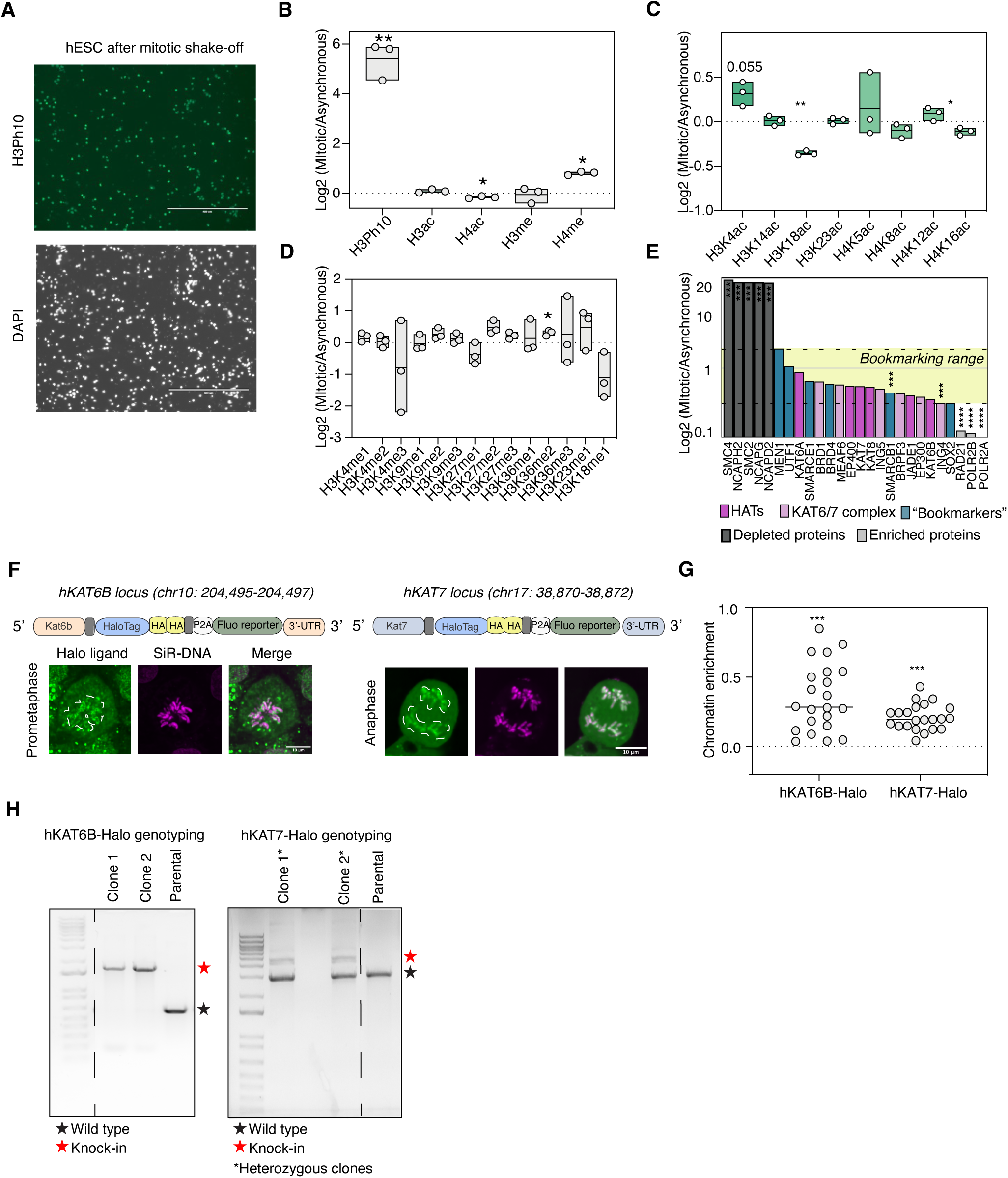
**A.** Representative immunofluorescence images showing the high purity of mitotic human ESCs after synchronization and shake-off. The mitotic marker, H3Ph10 (Histone-3-phosphoserine 10) is shown in green and DAPI in grey. **B.** Box plot showing the relative abundance of different histone modification marks on mitotic and asynchronous histone extracts (expressed as log_2_ ratio) from human ESCs as detected by mass-spectrometry. Data from n = 3 biological replicates is shown as a mean. Statistical significance was assessed by one-sample two-tailed t-test against 0. *p < 0.05, **p < 0.01. The full list of histone proteomics data is provided in Suppl. Table 1. **C.** Box plot comparing the relative abundance of select histone acetylation residues between mitotic and asynchronous extracts as log2 ratio. H3ac peptides are shown in dark green while H4ac peptides are light green. Data from n = 3 biological replicates are shown. Statistical significance was assessed by one-sample two-tailed t-test against 0. *p < 0.05, **p < 0.01. The full histone proteomics data is provided in Suppl. Table 1. **D.** Box plot showing widespread retention of different histone methylation marks on mitotic human ESCs relative to asynchronous extracts, as detected by mass-spectrometry. Data from n = 3 biological replicates is shown as a mean. Data from n = 3 biological replicates is shown as a mean. A statistically significant increase was only found for H3K36me2 upon one-sample two-tailed t-test against 0. * p < 0.05. The full list of histone proteomics data is provided in Suppl. Table 1. **E.** Bar plot of the mitotic/asynchronous ranking, log2foldchange(mitotic/asynchronous), in logarithmic scale for different categories of proteins. Proteins known to be either mitotically enriched or depleted are shown in dark gray or pale gray respectively. Previously described bookmarking factors in mESCs (since there are no bookmarking studies in hESCs) are shown in blue and are used to set the “bookmarking range” (yellow shadow limited by two dotted lines). Histone acetyltransferases (HATs), pink, and KAT6/7 complex subunits, violet, are present within the defined “bookmarking range”. n = 2 biological replicates per condition. Differential analysis was performed using a t-test. The full list of proteins for this figure is provided in Suppl. Table 2. **F.** *Top:* schematic of the HaloTag® CRISPR knock-in strategy in human ESCs to generate endogenous, fusion KAT6B-Halo and KAT7-Halo proteins for ligand-mediated live imaging. The knock-in cassettes were introduced into the endogenous *KAT7* and *KAT6B* gene *loci* after the last exon. In addition to the HaloTag® tag surrounded by linkers, the cassette contains 2xHA tags and a fluorescent transcriptional reporter preceded by P2A tag to facilitate cell-line generation. *Bottom:* live-cell imaging of mESCs HaloTagged for KAT6B and KAT7. Representative images show the HaloTagged protein in green in different mitotic phases. DNA is shown in magenta. Scale bar = 10 μm. **G.** Dot plot showing the chromatin enrichment (Chr. Enr) for both HaloTagged proteins calculated as log_2_(Chromosome intensity/Cytoplasmic intensity). Chr. Enr > 0.0 indicates mitotic enrichment while Chr. Enr < 0.0 shows relative depletion from mitotic chromosomes. n = 22 cells were quantified per protein. Both proteins were significantly enriched on mitotic chromatin as assessed by one-sample two-tailed t-test (***p< 0.001). **H.** Representative genotyping PCRs to verify successful knock-in of the HaloTag cassette at the endogenous *KAT6B* locus (homozygous) and *KAT7* locus (heterozygous) by CRISPR engineering compared to the wild-type bands in the parental lines. Primers are provided in Suppl. Table 7.

**Supp. Figure 6. Related to Figure 5.**
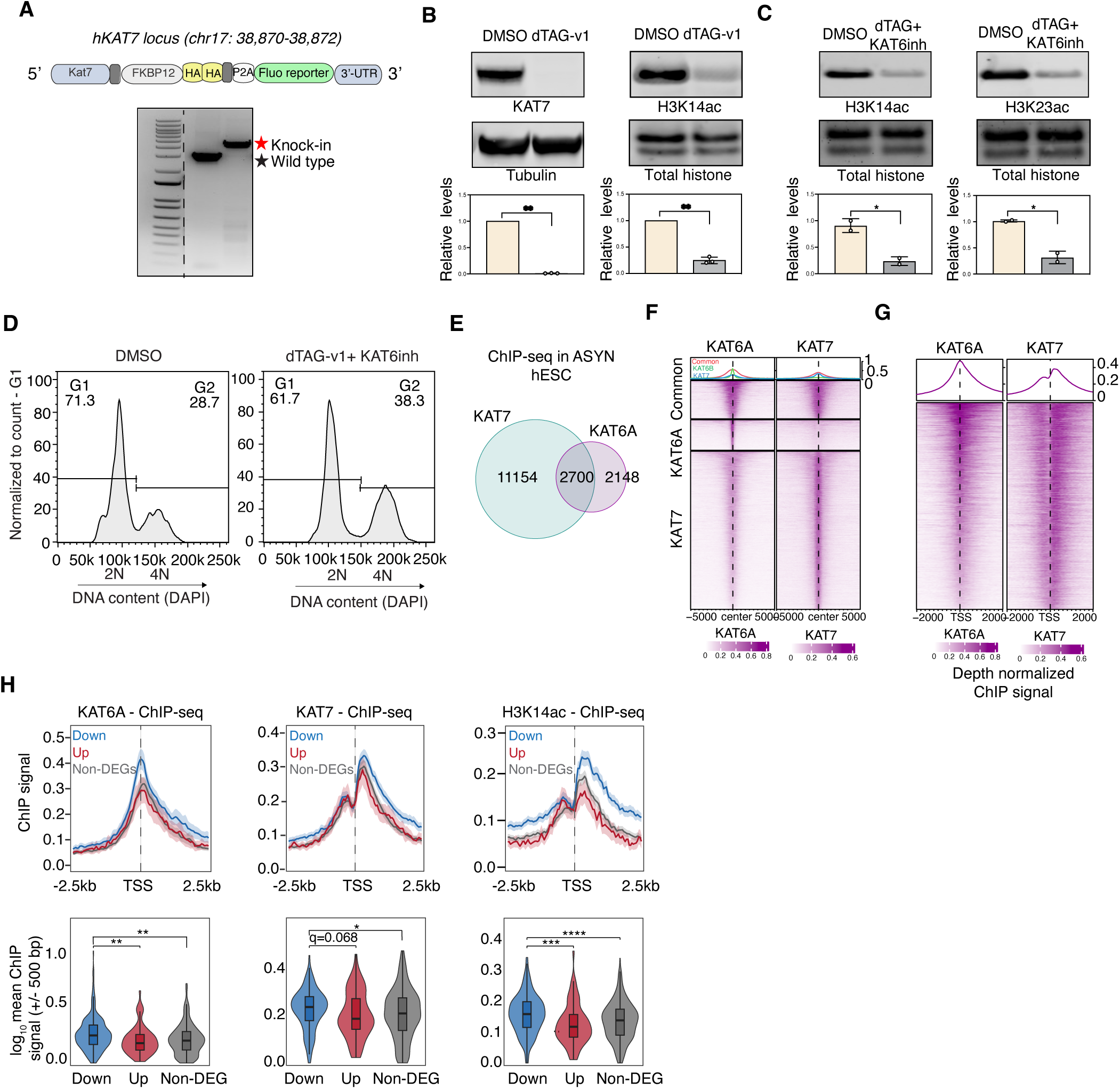
**A.** *Top:* Schematic of the CRISPR-mediated knock-in strategy to generate KAT7-degron hESCs. The knock-in cassette was introduced into the endogenous *KAT7 locus* after the last exon. hKAT7 is fused to the FKBP12-V, surrounded by linkers, 2xHA tags and a fluorescent transcriptional reporter preceded by P2A tag to facilitate detection and isolation of the successfully targeted cells. *Bottom:* Representative genotyping PCRs to verify successful homozygous knock-in of the FKBP12-V cassette at the endogenous *KAT7* locus compared to the wild-type band in the parental lines. Primers are provided in Suppl. Table 7. **B.** Validation of hKAT7 protein depletion and concomitant H3K14ac reduction in the homozygous hKAT7-FKBP12-V clones shown in panel (a) upon dTAG-v1 treatment. *Top:* Western blot analysis of KAT7 and H3K14ac levels after 48 h of DMSO or 250 nM dTAG-v1 treatment. Tubulin and total histone stain were used as loading controls respectively. *Bottom*: bar plot of the Western blot quantifications. Levels were normalized to the loading controls and plotted relative to DMSO controls. n = 3 biological replicates. KAT7 and H3K14ac levels were significantly reduced upon treatment as assessed by paired two-sided t-test (**p < 0.01). **C.** Western blot analysis (top) and quantification (bottom) of H3K14ac and H3K23ac levels in the hKAT7-FKBP12-V clones upon dual treatment with 250 nM dTAG-v1 and 100 nM KAT6 inhibitor for 2 hours prior to mitotic collection related to Fig.5A. Total histone stain was used as a loading control. Levels were normalized to the loading control and plotted relative to DMSO control. n = 2 technical replicates. H3K14ac and H3K23ac levels were significantly reduced upon treatment as assessed by paired two-sided t-test (*p < 0.05). **D.** FACS analysis measuring DAPI (DNA) content to quantify the successful release into G1 (for 5hr) of hESCs treated either with DMSO or with dual dTAG-V1/KAT6 inhibitor during mitotic arrest (**related to** Fig.5A). Cells showed a prominent 2N peak of similar magnitude in both control and treated cells indicating no cell-cycle defects in the release. **E.** Venn diagram showing the overlap for hKAT6A and hKAT7 ChIP-seq peaks in asynchronous hESCs. **F.** Tornado plot of the KAT6A and KAT7 ChIP-seq signal (normalized over input and sequencing depth) in asynchronous hESCs depicting common, KAT6A-specific and KAT7-specific peaks. The average signal of n= 2 independent biological replicates is projected +/- 5 kb around peak centers. **G.** Tornado plots showing the KAT6B and KAT7 ChIP-seq signals (normalized over input and sequencing depth) around their shared target genes. The average signal of n= 2 independent biological replicates is projected +/- 2kb around the Transcriptional Start Site (TSS). **H.** *Top:* metaplots showing the mean ChIP-seq signal for KAT6A, KAT7 and H3K14ac +/- 2.5kb around TSS of downregulated (blue), upregulated (red) and non-DEGs (grey) upon dual dTAG-V1 and KAT6i treatment in G1 hESCs, as described in Fig.5C. *Bottom:* Violin plots and statistical analysis comparing the mean ChIP-seq signal around +/- 500 bp TSS on the three gene categories. Downregulated genes had significantly higher mean ChIP-seq signal for KAT6A, KAT6 and H3K14ac compared to upregulated genes and non-DEGs as assessed by two sided Wilcoxon rank-sum test. *q < 0.05, **q < 0.01, ***q < 0.001, ****q < 0.0001.

**Supp. Figure 7. Related to Figure 5.**
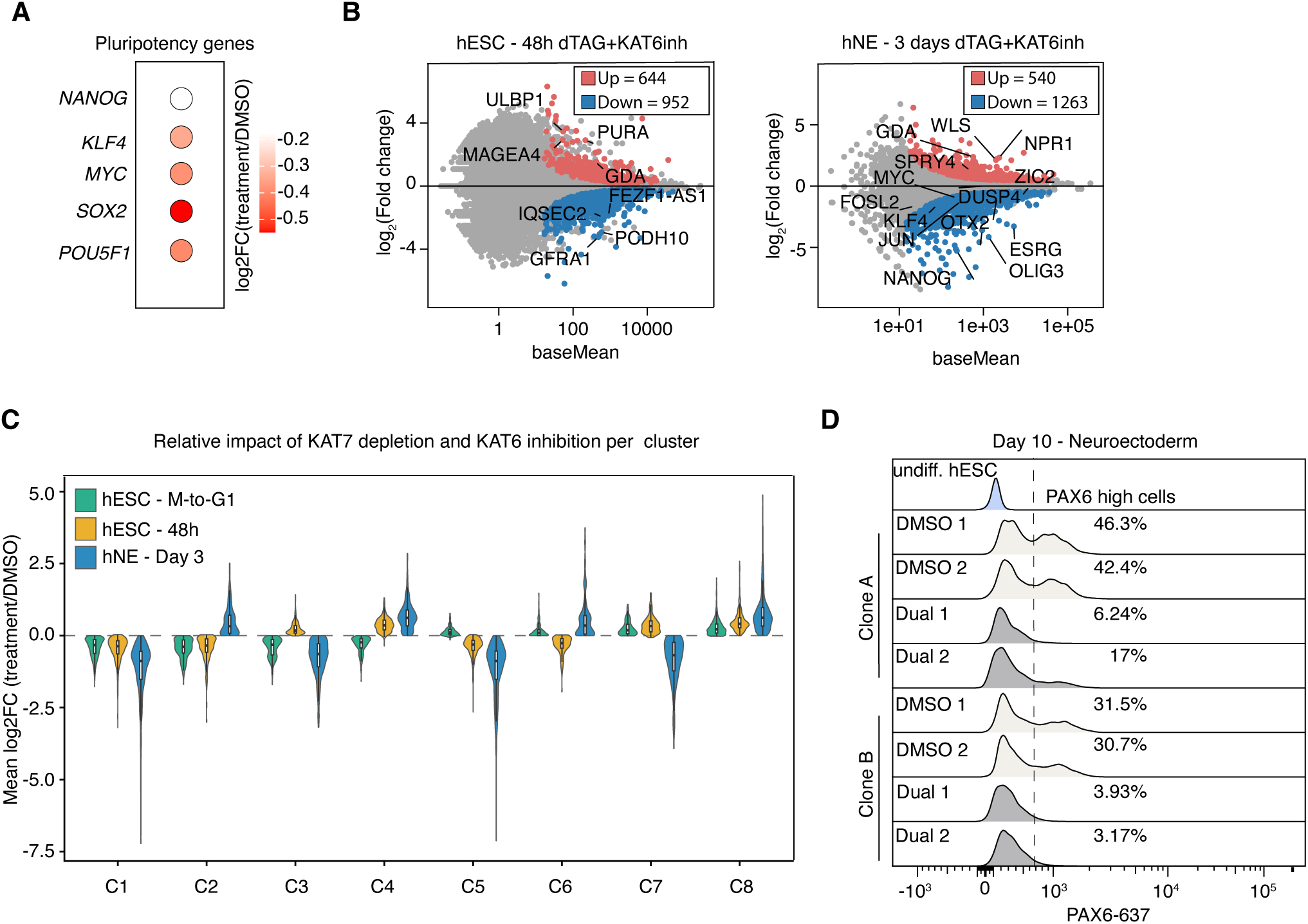
**A.** Dot plot showing the log2FC of core pluripotency factors in dTAG versus DMSO cells in G1. **B.** M/A plots of differentially expressed genes in DMSO versus treated (dTAG-v1+KAT6i) cells under different culture conditions; related to the heatmap in Fig. 5G. The log_2_ fold change (treated/DMSO) is plotted on the y-axis against the average expression for each gene (baseMean) on the x-axis for DMSO (n = 2 biological replicates). p-adj val > 0.05 was used as a cut-off to call differential expression. Each point represents a single gene. Up-regulated genes are colored in red and downregulated genes are colored in blue. Non-DEGs are grey. *Left:* M/A plot showing expression changes in hESC after a 48 h treatment. *Right:* M/A plot showing expression changes after dual treatment of hESCs undergoing neuroectoderm specification for 3 days. The full list of DEGs is provided in Suppl. Table 5. **C.** Violin plot showing the relative impact of KAT7 depletion and KAT6 inhibition per cluster in Fig.5G. The log2FC(treatment/DMSO) is shown for each condition. **D.** FACs analysis of PAX6 protein levels after intracellular staining in DMSO- or dTAG/KAT6I-treated hESCs after 10 days of neuroectoderm specification. Two independent clones with two technical replicates each were analyzed, while undifferentiated/untreated hESCs (blue) served as a negative control. The vertical dotted line marks the cut-off for the PAX6+ cells. The percentage of PAX6+ cells is depicted in Fig. 5J.

## References

1. Romito, A. & Cobellis, G. Pluripotent Stem Cells: Current Understanding and Future Directions. Stem Cells Int. 2016, (2016).

2. Evans, M. Discovering pluripotency: 30 years of mouse embryonic stem cells. Nat. Rev. Mol. Cell Biol. 12, 680–686 (2011).

3. Soufi, A. & Dalton, S. Cycling through developmental decisions: how cell cycle dynamics control pluripotency, differentiation and reprogramming. Development 143, 4301–4311 (2016).

4. Pauklin, S. & Vallier, L. The Cell-Cycle State of Stem Cells Determines Cell Fate Propensity. Cell 155, 135 (2013).

5. Dalton, S. Linking the Cell Cycle to Cell Fate Decisions. Trends Cell Biol. 25, 592–600 (2015).

6. Egli, D., Birkhoff, G. & Eggan, K. Mediators of reprogramming: Transcription factors and transitions through mitosis. Nat. Rev. Mol. Cell Biol. 9, 505–516 (2008).

7. Pelham-Webb, B., Murphy, D. & Apostolou, E. Dynamic 3D Chromatin Reorganization during Establishment and Maintenance of Pluripotency. Stem Cell Reports 15, 1176–1195 (2020).

8. Ito, K. & Zaret, K. S. Maintaining Transcriptional Specificity Through Mitosis. Annu. Rev. Genomics Hum. Genet. 23, 53–71 (2022).

9. Oomen, M. E. & Dekker, J. Epigenetic characteristics of the mitotic chromosome in 1D and 3D. Crit. Rev. Biochem. Mol. Biol. 52, 185–204 (2017).

10. Singh, A. M. et al. Cell-cycle control of developmentally regulated transcription factors accounts for heterogeneity in human pluripotent cells. Stem Cell Reports 1, 532–544 (2013).

11. Asenjo, H. G. et al. Polycomb regulation is coupled to cell cycle transition in pluripotent stem cells. Sci. Adv. 6, (2020).

12. Pelham-Webb, B. et al. H3K27ac bookmarking promotes rapid post-mitotic activation of the pluripotent stem cell program without impacting 3D chromatin reorganization. Mol. Cell 81, 1732–1748.e8 (2021).

13. Zaidi, S. K., Lian, J. B., van Wijnen, A., Stein, J. L. & Stein, G. S. Mitotic gene bookmarking: An epigenetic mechanism for coordination of lineage commitment, cell identity and cell growth. Adv. Exp. Med. Biol. 962, 95–102 (2017).

14. Kadauke, S. & Blobel, G. A. Mitotic bookmarking by transcription factors. Epigenetics Chromatin 6, 6 (2013).

15. Zaidi, S. K. et al. Mitotic gene bookmarking: An epigenetic program to maintain normal and cancer phenotypes. Molecular Cancer Research 16, 1617–1624 (2018).

16. Kang, H. et al. Dynamic regulation of histone modifications and long-range chromosomal interactions during postmitotic transcriptional reactivation. Genes Dev. 34, 913–930 (2020).

17. Kouskouti, A. & Talianidis, I. Histone modifications defining active genes persist after transcriptional and mitotic inactivation. EMBO J. 24, 347–357 (2005).

18. Blobel, G. A. et al. A Reconfigured Pattern of MLL Occupancy within Mitotic Chromatin Promotes Rapid Transcriptional Reactivation Following Mitotic Exit. Mol. Cell 36, 970–983 (2009).

19. Margueron, R. & Reinberg, D. Chromatin structure and the inheritance of epigenetic information. Nat. Rev. Genet. 11, 285–296 (2010).

20. Xu, M., Wang, W., Chen, S. & Zhu, B. A model for mitotic inheritance of histone lysine methylation. EMBO Rep. 13, 60–67 (2011).

21. Muramoto, T., Müller, I., Thomas, G., Melvin, A. & Chubb, J. R. Methylation of H3K4 Is required for inheritance of active transcriptional states. Curr. Biol. 20, 397–406 (2010).

22. Javasky, E. et al. Study of mitotic chromatin supports a model of bookmarking by histone modifications and reveals nucleosome deposition patterns. Genome Res. 28, 1455–1466 (2018).

23. Cimini, D., Mattiuzzo, M., Torosantucci, L. & Degrassi, F. Histone Hyperacetylation in Mitosis Prevents Sister Chromatid Separation and Produces Chromosome Segregation Defects. Mol. Biol. Cell 14, 3821 (2003).

24. Schneider, M. W. G. et al. A mitotic chromatin phase transition prevents perforation by microtubules. Nature 2022 609:7925 609, 183–190 (2022).

25. Turner, B. M. Acetylation and deacetylation of histone H4 continue through metaphase with depletion of more-acetylated isoforms and altered site usage. Exp. Cell Res. 182, 206–214 (1989).

26. Jeppesen, P., Mitchell, A., Turner, B. & Perry, P. Antibodies to defined histone epitopes reveal variations in chromatin conformation and underacetylation of centric heterochromatin in human metaphase chromosomes. Chromosoma 101, 322–332 (1992).

27. Zhao, R., Nakamura, T., Fu, Y., Lazar, Z. & Spector, D. L. Gene bookmarking accelerates the kinetics of post-mitotic transcriptional re-activation. Nat. Cell Biol. 13, 1295–1304 (2011).

28. Halsall, J. A. et al. Histone modifications form a cell-type-specific chromosomal bar code that persists through the cell cycle. Sci. Rep. 11, (2021).

29. Valis, E., Sánchez-Molina, S. & Martínez-Balbás, M. A. Role of histone modifications in marking and activating genes through mitosis. J. Biol. Chem. 280, 42592–42600 (2005).

30. Behera, V. et al. Interrogating Histone Acetylation and BRD4 as Mitotic Bookmarks of Transcription. Cell Rep. 27, 400–415.e5 (2019).

31. Kochanova, N. Y. et al. A time-resolved atlas of histone modifications during mitotic entry. Mol. Cell 86, 1653–1672.e9 (2026).

32. Liu, Y. et al. Widespread Mitotic Bookmarking by Histone Marks and Transcription Factors in Pluripotent Stem Cells. Cell Rep. 19, 1283–1293 (2017).

33. Hsiung, C. C.-S. et al. Genome accessibility is widely preserved and locally modulated during mitosis. Genome Res. 25, 213–225 (2015).

34. Turner, B. M. Histone acetylation and an epigenetic code. Bioessays 22, 836–845 (2000).

35. Patil, H. et al. Mitotic Activation of a Novel Histone Deacetylase 3-Linker Histone H1.3 Protein Complex by Protein Kinase CK2. J. Biol. Chem. 291, 3158 (2015).

36. Gandhi, S. et al. Mitotic H3K9ac is controlled by phase-specific activity of HDAC2, HDAC3, and SIRT1. Life Sci. Alliance 5, (2022).

37. Ginno, P. A., Burger, L., Seebacher, J., Iesmantavicius, V. & Schübeler, D. Cell cycle-resolved chromatin proteomics reveals the extent of mitotic preservation of the genomic regulatory landscape. Nat. Commun. 9, (2018).

38. Kruhlak, M. J. et al. Regulation of global acetylation in mitosis through loss of histone acetyltransferases and deacetylases from chromatin. J. Biol. Chem. 276, 38307–38319 (2001).

39. Djeghloul, D. et al. nature cell biology Hbo1 and Msl complexes preserve differential compaction and H3K27me3 marking of active and inactive X chromosomes during mitosis Check for updates. Nature Cell Biology | 27, 1482–1495 (2025).

40. Shvedunova, M. & Akhtar, A. Modulation of cellular processes by histone and non-histone protein acetylation. Nature Reviews Molecular Cell Biology vol. 23 329–349 Preprint at 10.1038/s41580-021-00441-y (2022).

41. Zheng, T., Wang, S., Liu, W. & Lu, Y. Targeting KAT6A/B as a New Therapeutic Strategy for Cancer Therapy. J. Med. Chem. 68, 1002–1020 (2025).

42. Gordon, S. J. V. et al. Catalytic Inhibition of KAT6/KAT7 Enhances the Efficacy and Overcomes Primary and Acquired Resistance to Menin Inhibitors in MLL Leukemia. Cancer Discov. 15, 2117–2138 (2025).

43. Michmerhuizen, N. L. et al. KAT6A and KAT7 Histone Acetyltransferase Complexes Are Molecular Dependencies and Therapeutic Targets in NUP98-Rearranged Acute Myeloid Leukemia. Cancer Discov. 15, 2096–2116 (2025).

44. Kueh, A. J. et al. Stem cell plasticity, acetylation of H3K14, and de novo gene activation rely on KAT7. Cell Rep. 42, 111980 (2023).

45. Voss, A. K. et al. KAT6A is essential for developmental control gene expression in neural stem and progenitor cells. PLoS Genet. 22, e1012133 (2026).

46. Bergamasco, M. I. et al. Increasing histone acetylation improves sociability and restores learning and memory in KAT6B-haploinsufficient mice. J. Clin. Invest. 134, (2024).

47. Clayton-Smith, J. et al. Whole-exome-sequencing identifies mutations in histone acetyltransferase gene KAT6B in individuals with the Say-Barber-Biesecker variant of Ohdo syndrome. Am. J. Hum. Genet. 89, 675–681 (2011).

48. Day, R. et al. A clinical and genetic study of the Say/Barber/Biesecker/Young-Simpson type of Ohdo syndrome. Clin. Genet. 74, 434–444 (2008).

49. Tham, E. et al. Dominant mutations in KAT6A cause intellectual disability with recognizable syndromic features. Am. J. Hum. Genet. 96, 507–513 (2015).

50. Arboleda, V. A. et al. De Novo Nonsense Mutations in KAT6A, a Lysine Acetyl-Transferase Gene, Cause a Syndrome Including Microcephaly and Global Developmental Delay. Am. J. Hum. Genet. 96, 498 (2015).

51. Sidoli, S. et al. One minute analysis of 200 histone posttranslational modifications by direct injection mass spectrometry. Genome Res. 29, 978– 987 (2019).

52. Zheng, Y., Thomas, P. M. & Kelleher, N. L. Measurement of acetylation turnover at distinct lysines in human histones identifies long-lived acetylation sites. Nat. Commun. 4, (2013).

53. Zaret, K. S. Genome reactivation after the silence in mitosis: Recapitulating mechanisms of development? Dev. Cell 29, 132–134 (2014).

54. Zhu, Z. et al. Mitotic bookmarking by SWI/SNF subunits. Nature 618, 180– 187 (2023).

55. Djeghloul, D. et al. Identifying proteins bound to native mitotic ESC chromosomes reveals chromatin repressors are important for compaction. Nat. Commun. 11, 4118 (2020).

56. Hsiung, C. C.-S. et al. A hyperactive transcriptional state marks genome reactivation at the mitosis–G1 transition. Genes Dev. 30, 1423–1439 (2016).

57. Festuccia, N. et al. Mitotic binding of Esrrb marks key regulatory regions of the pluripotency network. Nat. Cell Biol. 18, 1139–1148 (2016).

58. Shen, Y. et al. TBP bookmarks and preserves neural stem cell fate memory by orchestrating local chromatin architecture. Mol. Cell 85, 413–429.e10 (2025).

59. Xing, H., Vanderford, N. L. & Sarge, K. D. The TBP-PP2A mitotic complex bookmarks genes by preventing condensin action. Nat. Cell Biol. 10, 1318– 1323 (2008).

60. Nabet, B. et al. The dTAG system for immediate and target-specific protein degradation. Nat. Chem. Biol. 14, 431–441 (2018).

61. Bergamasco, M. I. et al. KAT6B is required for histone 3 lysine 9 acetylation and SOX gene expression in the developing brain. Life Sci. Alliance 8, (2025).

62. Christova, R. & Oelgeschläger, T. Association of human TFIID-promoter complexes with silenced mitotic chromatin in vivo. Nat. Cell Biol. 4, 79–82 (2002).

63. Seneviratne, J. A., Ho, W. W. H., Glancy, E. & Eckersley-Maslin, M. A. A low-input high resolution sequential chromatin immunoprecipitation method captures genome-wide dynamics of bivalent chromatin. Epigenetics Chromatin 17, 3-(2024).

64. Bernstein, B. E. et al. A bivalent chromatin structure marks key developmental genes in embryonic stem cells. Cell 125, 315–326 (2006).

65. Karmodiya, K., Krebs, A. R., Oulad-Abdelghani, M., Kimura, H. & Tora, L. H3K9 and H3K14 acetylation co-occur at many gene regulatory elements, while H3K14ac marks a subset of inactive inducible promoters in mouse embryonic stem cells. BMC Genomics 13, 424 (2012).

66. Sharma, S. et al. Discovery of a highly potent, selective, orally bioavailable inhibitor of KAT6A/B histone acetyltransferases with efficacy against KAT6A-high ER+ breast cancer. Cell Chem. Biol. 30, 1191–1210.e20 (2023).

67. Ciceri, G. et al. An epigenetic barrier sets the timing of human neuronal maturation. Nature 2024 626:8000 626, 881–890 (2024).

68. Xu, N. et al. Genome-wide CRISPR screen identifies Menin and SUZ12 as regulators of human developmental timing. Nature Cell Biology 2025 27:9 27, 1411–1421 (2025).

69. Banerjee-Basu, S. & Packer, A. SFARI Gene: an evolving database for the autism research community. Dis. Model. Mech. 3, 133–135 (2010).

70. Lodhi, N., Ji, Y. & Tulin, A. Mitotic bookmarking: maintaining post-mitotic reprogramming of transcription reactivation. Curr. Mol. Biol. Rep. 2, 10 (2016).

71. Biggs, R., Liu, P. Z., Stephens, A. D. & Marko, J. F. Effects of altering histone posttranslational modifications on mitotic chromosome structure and mechanics. Mol. Biol. Cell 30, 820 (2019).

72. Valis, E., Sánchez-Molina, S. & Martínez-Balbás, M. A. Role of histone modifications in marking and activating genes through mitosis. J. Biol. Chem. 280, 42592–42600 (2005).

73. Djeghloul, D. et al. Loss of H3K9 trimethylation alters chromosome compaction and transcription factor retention during mitosis. Nat. Struct. Mol. Biol. 30, 489–501 (2023).

74. Bryan, E. et al. Nucleosomal asymmetry shapes histone mark binding and promotes poising at bivalent domains. Mol. Cell 85, 471–489.e12 (2025).

75. MacPherson, L. et al. HBO1 is required for the maintenance of leukaemia stem cells. Nature 2019 577:7789 577, 266–270 (2019).

76. Yu, X. et al. An EGFR co-amplified lncRNA HELDR promotes glioblastoma malignancy through KAT7-driven gene programs. Nature Cell Biology 2026 28:5 28, 972–985 (2026).

77. Wang, H. et al. Histone acetylation by HBO1 (KAT7) activates Wnt/β-catenin signaling to promote leukemogenesis in B-cell acute lymphoblastic leukemia. Cell Death Dis. 14, 498 (2023).

78. Tan, Y., Wang, J. & Ma, F. Lysine Acetyltransferase 6 in Health and Disease. MedComm (Beijing). 6, e70520 (2025).

79. Mukohara, T. et al. A phase 1 dose expansion study of a first-in-class KAT6 inhibitor (PF-07248144) in patients with advanced or metastatic ER+ HER2− breast cancer. Journal of Clinical Oncology 42, (2024).

80. Mukohara, T. et al. Inhibition of lysine acetyltransferase KAT6 in ER+HER2− metastatic breast cancer: a phase 1 trial. Nature Medicine 2024 30:8 30, 2242–2250 (2024).

81. Gupta, S. et al. MAGMa: Your Comprehensive Tool for Differential Expression Analysis in Mass-Spectrometry Proteomic Data. *bioRxiv* 2024.06.24.600424 (2024) doi:10.1101/2024.06.24.600424.

82. Hsu, C. C. et al. Gas41 links histone acetylation to H2A.Z deposition and maintenance of embryonic stem cell identity. Cell Discovery 2018 4:1 4, 28-(2018).

83. Sheffield, N. C. & Bock, C. LOLA: enrichment analysis for genomic region sets and regulatory elements in R and Bioconductor. Bioinformatics 32, 587– 589 (2016).

84. Ee, L. sha et al. Enhancer remodeling by OTX2 directs specification and patterning of mammalian definitive endoderm. Dev. Cell 60, (2025).

85. Lee, U. et al. Post-mitotic transcriptional activation and 3D regulatory interactions show locus-and di9erentiation-specific sensitivity to cohesin depletion 2 3. 10.1101/2025.02.13.638153 doi:10.1101/2025.02.13.638153.

86. Sidoli, S., Bhanu, N. V., Karch, K. R., Wang, X. & Garcia, B. A. Complete Workflow for Analysis of Histone Post-translational Modifications Using Bottom-up Mass Spectrometry: From Histone Extraction to Data Analysis. J. Vis. Exp. 2016, 54112 (2016).

87. Yuan, Z. F. et al. EpiProfile 2.0: A Computational Platform for Processing Epi-Proteomics Mass Spectrometry Data. J. Proteome Res. 17, 2533–2541 (2018).

88. Teves, S. S. et al. A dynamic mode of mitotic bookmarking by transcription factors. Elife 5, (2016).

89. Chervova, A. et al. Mitotic bookmarking redundancy by nuclear receptors in pluripotent cells. Nature Structural & Molecular Biology 2024 31:3 31, 513–522 (2024).

90. Buenrostro, J. D., Wu, B., Chang, H. Y. & Greenleaf, W. J. ATAC-seq: A Method for Assaying Chromatin Accessibility Genome-Wide. Curr. Protoc. Mol. Biol. 109, 21.29.1-21.29.9 (2015).

91. Köster, J. & Rahmann, S. Snakemake—a scalable bioinformatics workflow engine. Bioinformatics 28, 2520–2522 (2012).

92. Love, M. I., Huber, W. & Anders, S. Moderated estimation of fold change and dispersion for RNA-seq data with DESeq2. Genome Biol. 15, (2014).

93. Quinlan, A. R. & Hall, I. M. BEDTools: a flexible suite of utilities for comparing genomic features. Bioinformatics 26, 841–842 (2010).

94. Langmead, B. & Salzberg, S. L. Fast gapped-read alignment with Bowtie 2. Nat. Methods 9, 357–359 (2012).

95. Yu, G., Wang, L. G., Han, Y. & He, Q. Y. clusterProfiler: an R package for comparing biological themes among gene clusters. OMICS 16, 284–287 (2012).

96. Ritchie, M. E. et al. limma powers differential expression analyses for RNA-sequencing and microarray studies. Nucleic Acids Res. 43, e47–e47 (2015).

97. Huang, T. et al. MSstatsTMT: Statistical Detection of Differentially Abundant Proteins in Experiments with Isobaric Labeling and Multiple Mixtures. Molecular and Cellular Proteomics 19, 1706–1723 (2020).

98. Zhou, Y. et al. A comprehensive SARS-CoV-2–human protein–protein interactome reveals COVID-19 pathobiology and potential host therapeutic targets. Nature Biotechnology 2022 41:1 41, 128–139 (2022).

99. Perez-Riverol, Y. et al. The PRIDE database at 20 years: 2025 update. Nucleic Acids Res. 53, D543–D553 (2025).

